# Flexible neural representations of abstract structural knowledge in the human Entorhinal Cortex

**DOI:** 10.1101/2023.08.31.555760

**Authors:** S. Mark, P. Schwartenbeck, A Hahamy, V Samborska, AB. Baram, TEJ Behrens

**Author notes:** Equal contribution (AB. Baram, TEJ Behrens).

## Abstract

Humans’ ability for generalisation is outstanding. It is flexible enough to identify cases where knowledge from prior tasks is relevant, even when many features of the current task are different, such as the sensory stimuli or the size of the task state space. We have previously shown that in abstract tasks, humans can generalise knowledge in cases where the only cross-task shared feature is the statistical rules that govern the task’s state-state relationships. Here, we hypothesized that this capacity is associated with generalisable representations in the entorhinal cortex (EC). This hypothesis was based on the EC’s generalisable representations in spatial tasks and recent discoveries about its role in the representation of abstract tasks. We first develop an analysis method capable of testing for such representations in fMRI data, explain why other common methods would have failed for our task, and validate our method through a combination of electrophysiological data analysis, simulations and fMRI sanity checks. We then show with fMRI that EC representations generalise across complex non-spatial tasks that share a hexagonal grid structural form but differ in their size and sensory stimuli, i.e. their only shared feature is the rules governing their statistical structure. There was no clear evidence for such generalisation in EC for non-spatial tasks with clustered, as opposed to planar, structure.

## Introduction

If you grew up in a small town, arriving in a big city might come as a shock. However, you’ll still be able to make use of your previous experiences, despite the difference in the size of the environment: When trying to navigate the busy city streets, your knowledge of navigation in your hometown is crucial. For example, it’s useful to know the constraints that a 2D topological structure exerted on distances between locations. When trying to make new friends, it’s useful to remember how people in your hometown tended to cluster in groups, with popular individuals perhaps belonging to several groups. Indeed, the statistical rules (termed “structural form”, (Kemp and Tenenbaum 2008)) that govern the relationships between elements (states) in the environment are particularly useful for generalisation to novel situations, as they do not depend on the size, shape or sensory details of the environment (Mark *et al*. 2020). Such generalisable features of environments are proposed to be part of the “cognitive map” encoding the relationships between their elements (Tolman 1948; Behrens *et al*. 2018; Mark *et al*. 2020).

The most studied examples of such environments are spatial 2D tasks. In all spatial environments, regardless of their size or shape, the relations between states (in this case locations) are subject to the same Euclidean statistical constraints. The spatial example is particularly useful because neural spatial representations are well-characterised. Indeed, one of the most celebrated of these - grid cells in the entorhinal cortex (EC) - has been suggested as (part of) a neural substrate for spatial generalisation (Behrens *et al*. 2018; Whittington *et al*. 2022). This is because (within a grid module) grid cells maintain their coactivation structure across different spatial environments (Fyhn *et al*. 2007; Yoon *et al*. 2013). In other words, the information embedded in grid cells generalises across 2D spatial environments (including environments of different shapes and sizes). Following a surge of studies showing that EC spatial coding principles are also used in non-spatial domains (Constantinescu, O’Reilly and Behrens 2016; Garvert, Dolan and Behrens 2017; Bao *et al*. 2019; Park *et al*. 2020), we have recently shown that EC also generalises over non-spatial environments that share the same statistical structure (Baram *et al*. 2021). Importantly, in that work the graphs that described the same-structured environments were isomorphic - i.e. there was a one-to-one mapping between states across same-structure environments.

What do we mean when we say the EC has “generalisable representations” in spatial tasks? and how can we probe these representations in complex non-spatial tasks? Between different spatial environments, each grid cell realigns: its firing fields might rotate and shift (Fyhn *et al*. 2007). Crucially, this realignment is synchronized within a grid module population (Yoon *et al*. 2013; Gardner *et al*. 2022), such that the change in the grid angle and phase of all cells is the same. This means that cells that have neighboring firing fields in one environment will also have neighboring firing fields in another environment-the coactivation structure is maintained (Yoon *et al*. 2013; Gardner *et al*. 2022). A mathematical corollary is that grid cells’ activity lies in the same low-dimensional subspace (manifold, (Yoon *et al*. 2013; Gardner *et al*. 2022)) in all spatial environments. This subspace remains even during sleep, meaning the representation is stably encoded (Burak and Fiete 2009; Gardner *et al*. 2019; Trettel SG *et al*. 2019).

We have recently developed an analysis method, referred to as “subspace generalisation”, which allows for the quantification of the similarities between linear neural subspaces, and used it to probe generalisation in cell data (Samborska *et al*. 2022). Unlike other representational methods for quantifying the similarity between activity patterns (like RSA, used in Baram *et al*. (Kriegeskorte, Mur and Bandettini 2008; Diedrichsen and Kriegeskorte 2017)), this method has the ability to isolate the shared features underlying tasks that do not necessarily have a straightforward cross-task mapping between states, such as when the sizes of tasks underlying graphs are different. Here, we use it to quantify generalisation in such a case, but on fMRI data of humans solving complex abstract tasks rather than on cell data. We designed an abstract associative-learning task in which visual images were assigned to nodes on a graph and were presented sequentially, according to their relative ordering on the graph. The graphs belonged to two different families of graphs, each governed by a different set of statistical regularity rules (structural forms (Kemp and Tenenbaum 2008)) – hexagonal (triangular) lattice graphs, and community structure graphs. There were two graphs of each structural form. Crucially, the graph size and embedded images differed within a pair of graphs with the same structural form (Figure 3b), allowing us to test generalisation due to structural form across both environment size and sensory information.

We first validate our approach by showing that subspace generalisation detects the known generalisation properties of entorhinal grid cells and hippocampal place cells when rodents free-forage in two different spatial environments – properties that have inspired our study’s hypothesis. Next, we propose that our method can capture these properties even in low-resolution data such as fMRI. We provide twofold support for this conjecture: through sampling and averaging of the rodent data to create low resolution version of the data, and through simulations of grid cells grouped into simulated voxels to account for the very low resolution of the BOLD signal. We use these simulations to discuss how the sensitivity of our method depends on various characteristics of the signal. Next, we validate the method for real fMRI signals by showing it detects known properties of visual encoding in the visual cortex in our task. Finally, and most importantly, we show that EC generalises its voxelwise covariance patterns over abstract, discrete hexagonal graphs of different size and stimuli, exactly as grid cells do in space. This result, however, did not hold for the community graph structures. We discuss some possible experimental shortcomings that might have led to this null result.

### Theory – “subspace generalisation”

How can we probe the neural correlates of generalisation of abstract tasks in the human brain? Popular representational analysis methods such as Representational Similarity Analysis (RSA) (Kriegeskorte, Mur and Bandettini 2008; Diedrichsen and Kriegeskorte 2017) and Repetition Suppression (Grill-Spector, Henson and Martin 2006; Barron, Garvert and Behrens 2016) have afforded some opportunities in this respect (Baram *et al*. 2021). However, because these methods rely on similarity measures between task states, they require labeling of a hypothesized similarity between each pair of states across tasks. Such labeling is not possible when we do not know which states in one task align with which states in another task. In the spatial example where states are locations, the mapping of each location in room A to locations in room B doesn’t necessarily exist - particularly when the rooms differ in size or shape. This makes labeling of hypothesized similarity between each pair of locations impossible. How can we look for shared activity patterns in such a case?

We have recently proposed this can be achieved by studying the covariance of different neurons across states (Samborska *et al*. 2022) (as opposed to RSA - which relies on the correlation of different states across neurons). If two tasks contain similar patterns of neural activity (regardless of when these occurred in each task), then the *neuron X neuron* covariance matrix (across states within-task) will look similar in both tasks. This covariance matrix can be summarised by its principal components (PCs), which are patterns across neurons - akin to “cell assemblies” - and their eigenvalues, which indicate how much each pattern contributes to the overall variance in the data. If representations generalise across tasks, then patterns that explain a lot of variance in task 1 will also explain a lot of variance in task 2. We can compute the task 2 variance explained by each of the PCs of task 1:

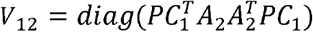

Where PC_1_ is a matrix with all task 1 PCs as its columns, ordered by their eigenvalues, and A_2_ is the *neurons X states* task 2 data. These PCs are ordered according to the variance explained in task 1. Hence, if the same PCs explain variance across tasks, early PCs will explain more variance in task 2 than late PCs. The cumulative sum of *V*_12_ will be a concave function and the area under this concave function is a measure of how well neuronal patterns generalise across tasks (Figure 1a). We refer to this measure as subspace generalisation.

**Figure 1.**
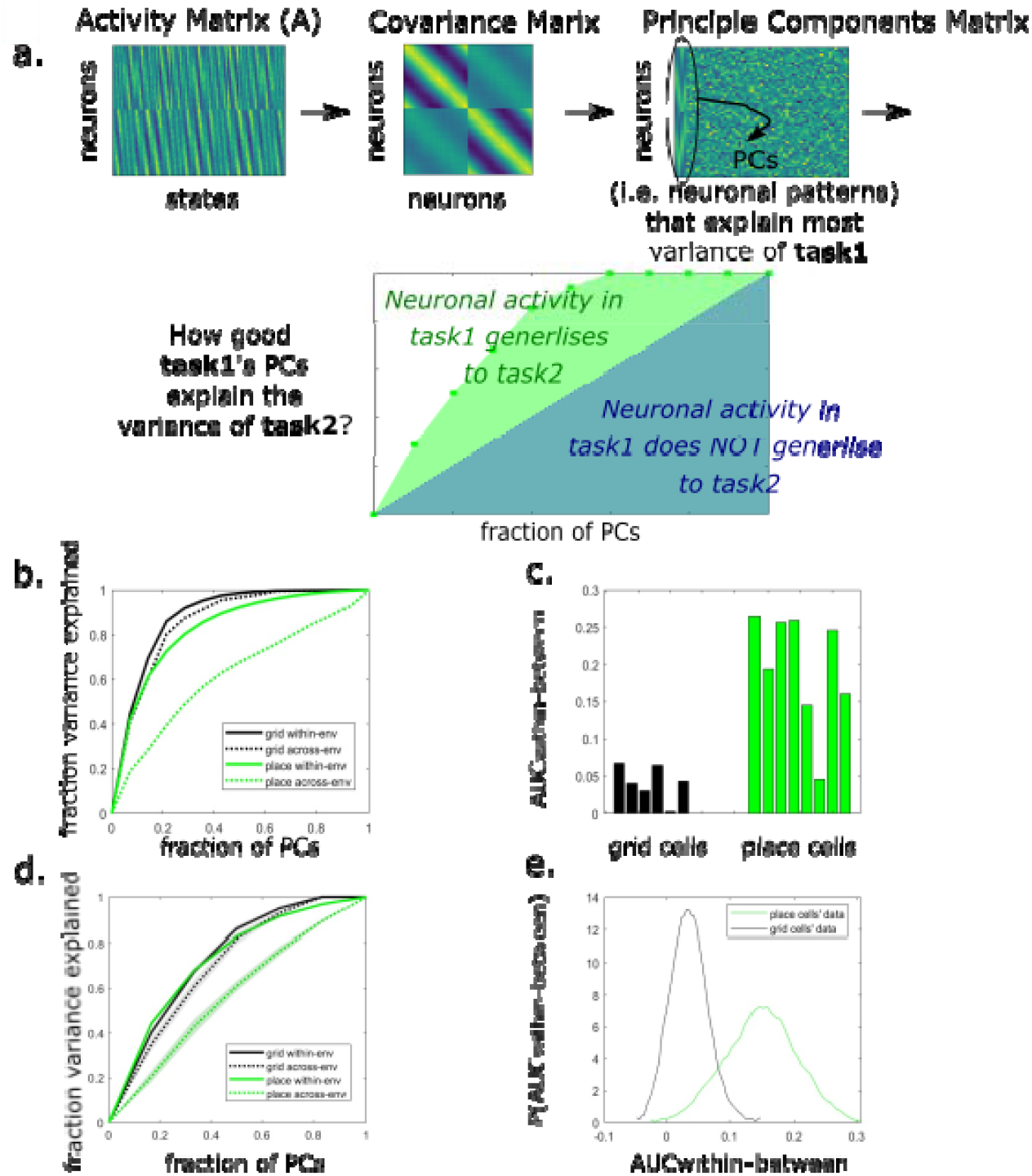
Subspace generalisation across environments in grid and place cells in data from Chen et al. 2018. a. Illustration of the subspace generalization method. The Principal Components (PCs) are calculated using the covariance matrix of the neuronal activity matrix. Then the activity matrix is projected on each PC (recorded when the animal was in the same or different environment/task) and the variance explained along each PC dimension is calculated. We calculate the Area Under the Curve (AUC) of the cumulative sum of the variance explained on each PC’s dimension as our similarity measure. When the similarity in neuronal patterns during the two different tasks is higher the area under the curve is higher (green AUC is added to the blue AUC) b. The cumulative variance explained by the PCs calculated using the activity of grid (black) or place (green) cells, within (solid lines) and across (dotted lines) environments. Subspace generalization is calculated as the difference between the area under the curve (AUC) of two lines. The difference between the black lines is small, indicating generalisation of grid cells across environments. The difference between the green lines is larger, indicating remapping of place cells (p<0.001, permutation test, see Methods). c. The difference between the within and across (solid and dashed lines in a., respectively) environments AUCs of the cumulative variance explained by grid or place cells (black or green lines in a., respectively). Data shown for all mice with enough grid or place cells (>10 recorded cells of the same type, each bar is a mouse and a specific projection (i.e. projecting on environment one or two)). The differences between the grid cells AUCs are significantly smaller than the place cells (p < 0.001 permutation test, see supplementary for more statistical analyses and specific examples). d. An example of the cumulative variance explained by the PCs, calculated using the constructed low-resolution version of grid and place cells data. The solid and dotted lines are average over 10 samples and the shaded areas represent the standard error of the mean across samples. Here, as above, the solid lines are projection within environment and the dotted lines are projections between environments. e. Subspace generalization in the low resolution version of the data captures the same generalization properties of grid vs place cells. The distributions were created via bootstrapping over cells from the same animal, averaging their activity, concatenating the samples across all animals and calculating the AUC difference between within and across environments projections (p<<0.001 Kolmogorov-Smirnovtest).

As validation and demonstration of our method, we first use it to recover differences in generalisation between grid cells and place cells in the rodent brain that have been shown previously with other methods. Next, we demonstrate the feasibility of our method in capturing this difference in generalization properties even after we manipulate the data and reduce its resolution. To complete the logical bridge from cells to voxels, we address the limitation of this demonstration: the low number of cells recorded. We simulate voxels from synthetic grid cells and show how our method’s power depends on various characteristics of the signal. These analyses show that theoretically (and under reasonable conditions) our method could still detect medial temporal lobe generalisation properties in fMRI BOLD signal. Finally, and most importantly, we use our method to analyse fMRI data, testing for generalisation of the covariance between voxel representations in human EC across complex non-spatial graphs with common regularities – analogous to the generalisation of grid cells in physical space. Crucially, in this task other representational methods common in fMRI analysis such as RSA or repetition suppression would not be applicable (due to lack of one-to-one mapping between states across graphs), highlighting the usefulness of our method.

## Results

### Subspace generalization captures known generalisation properties of grid and place cells

Grid cells and place cells differ in their generalisation property. When an animal moves from one environment to another, place cells “remap”: they change their correlation structure such that place cells that are neighbours in environment 1 need not be neighbors in environment 2. By contrast grid cells do not remap: the correlation structure between grid cells is preserved across environments, such that pairs of grid cells (within the same module) that have neighboring fields in environment 1 will also have neighboring fields in environment 2 (Fyhn *et al*. 2007). This is true even though each grid cell shifts and rotates its firing fields across environments - the grid cell population within a module realigns in unison (Gardner *et al*. 2022; Waaga *et al*. 2022). Crucially, the angle and phase of this realignment can’t be predicted in advance, meaning it is not possible to create hypotheses to test regarding the similarity between representations at a given location in environment 1 and a given location in environment 2 - a requirement for fMRI-compatible methods such as RSA or repetition suppression. In this section we demonstrate how subspace generalisation - which can also be useful in fMRI - captures the generalisation properties of grid and place cells that have previously been shown only with traditional analysis methods that require access to firing maps of single cells.

We computed subspace generalisation for grid and place cells recorded with electrophysiology in a previous study (Chen *et al*. 2018), in which mice freely-foraged in two square environments: a real physical and a virtual reality (VR) (see Methods for more details). For our purposes, this dataset is useful because large numbers of both place cells and grid cells were recorded (concurrently within a cell type) in two different environments - rather than because of the use of a VR environment.

We compared two different situations: one where “task 1” and “task 2” were actually from the same environment, Figure 1a - solid line, within-environment) and one where “task 1” and “task 2” were from different environments (Figure 1a - dotted line, across-environments).

As predicted, across environments grid cells’ subspaces generalised: PCs that were calculated using activity in one environment explained the activity variance in the other environment just as well as the within-environment baseline (Figure 1a, compare dotted and solid black lines, plots show the average of the projections of activity from one environment on EVs from the other environment and vice versa). The difference between the area under the curve (AUC) of the two lines was significantly smaller than chance (p<0.001 using a permutation test, see Methods and supplementary Figure S1). Importantly, grid cells generalized much better between the environments than place cells; the difference in AUCs between the solid and dotted lines is significantly smaller for grid cells compared to place cells (Figure 1b, p<0.001, for both permutation test and 2 sample t-test, see Methods and supplementary material). Interestingly, the difference in AUCs was also significantly smaller than chance for place cells (Figure 1a, compare dotted and solid green lines, p<0.05 using permutation tests, see statistics and further examples in supplementary material Figure S2), consistent with recent models predicting hippocampal remapping that is not fully random (Whittington *et al*. 2020).

### From neurons to voxels

So far, we have validated our method when applied to neurons. However, our primary interest in this manuscript is to apply it to fMRI data. To illustrate the efficacy of this approach in revealing generalisable neuronal subspaces within low resolution data like fMRI, we applied our method to such data – both from manipulated electrophysiology and simulations. We first examined our method on low-resolution versions of the Chen *et al*. rodent MTL data, obtained by grouping and averaging cells. We show that our method can still detect subspace generalization even on the supra-cellular level. However, due to the small number of recorded cells, this analysis does not fully replicate a voxel’s BOLD signal, which corresponds to the average activity of thousands of cells. To address this, we simulated many grid cells and grouped them into voxels, with each voxel’s activity corresponding to the average activity of its cells. We then applied subspace generalisation to the simulated pseudo-voxels, and examined how the results depend on various signal characteristics.

Using Chen *et al* electrophysiology dataset, we first normalised each cell’s firing rate maps, and then created bootstrapped low-resolution data: for each sampling iteration we sampled 7 cells (with repeats) into 2 groups within each animal and averaged the activities of cells within each group. This results in a 2-long vector for each animal. We then concatenate these vectors across animals. Note that for grid cells, this pooling over independent groups of neurons is reminiscent of pooling over different grid modules in a single subject. For each sample we calculated the difference in the area under the curve (AUC) between within and across environments projections as above (averaged over the projections on both environments, Figure 1c). We repeat this bootstrapping step to create a distribution of the differences in AUC for place cells and grid cells (Figure 1d). The difference in AUC was smaller for grid cells than for place cells (p<0.001 Kolmogorov-Smirnovtest), as is expected from the single cells’ analysis above.

The required number of cells to simulate a voxel’s activity (l*et al*one multiple voxels) far exceeds the number of cells in the Chen *et al*. dataset. To overcome this limitation and support our conjecture that our method can detect subspace-generalization even in fMRI BOLD signal, we next used simulated data. We simulated grid cells (see methods) organized into four grid modules, each composed of more than 10000 cells. We organized the cells in each module into four groups (pseudo-voxels) and averaged the activity within each group (see supplementary info for an example of our analysis using different number of groups within each module, and how our results are affected by the number of voxels per module, Figure S3). We concatenated the pseudo-voxels from all modules into one vector and calculated the difference in subspace-generalization measure (i.e. the AUC of within and between environments). We explored how two characteristics of the data affect subspace generalization: whether the grouping into voxels (within each module) was organized according to grid phase, and the level of noise in the data.

We first grouped the cells into voxels randomly, i.e. without any a-priori assumption on the relationship between the physical proximity of cells within the cortical layer and their firing rate maps. Examples of the resulted “pseudo-voxels” activity maps can be seen in Figure 2a. However, recent work has suggested there is a relationship between grid cells’ physical proximity and their grid phases (Gu *et al*. 2018). We therefore also simulated “pseudo voxels” by grouping grid cells, within each module, according to their grid phase (Figure 2b). The pseudo-voxel’s signal in the latter case is substantially stronger (compare color bar scales a between Figure 2a and 2b).

**Figure 2:**
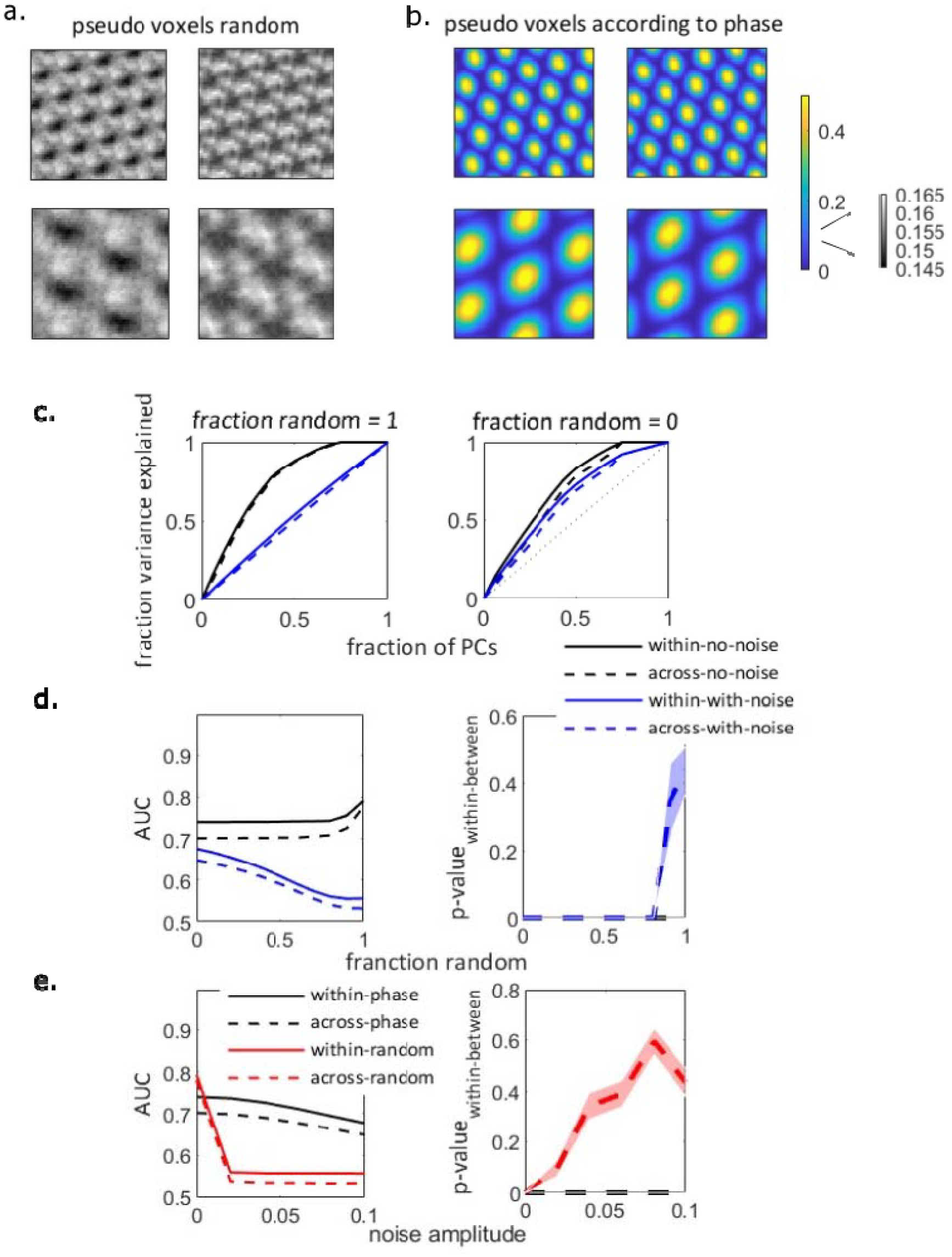
simulated voxels from simulated grid modules. a. Examples of simulated voxels activity map in the two environments, without noise. upper: higher frequency module, lower: lower frequency module. Cells are grouped into voxels randomly. b. Same as a. but with cells grouped into voxels according to the grid phase. Note the different scale of the color-bar between a. and b. c. Subspace generalization plot for the 16 simulated voxels, where the grouping into voxels is either random (left) or according to phase (right). Legend as in d, noise amplitude = 0.1. d. Left: AUCs of the subspace generalisation plots in c. as a function of the ratio of random vs phase-organised cells in the voxels, with no noise (black) or with high amplitude of noise (blue, *noise amplitude* = *0*.*1*). Without noise (black lines), the subspace generalization measure (AUC) remains high even when the fraction of randomly sampled cells increases. However, in the presence of noise, the subspace generalization measure decreases with the fraction of randomly sampled cells. Right: p-value of the effect according to the permutation distribution (see methods, shaded area: standard error of the mean). In the presence of noise and when the cells are sampled randomly, *AUC*_*within-between*_ becomes non-significant, see supplementary info Figure S3 for the dependency of the permutation distributions on the presence of noise and sampling. e. Same as d., except the continuous X-axis variable is the noise amplitude, for either of phase-organized (black) or randomly organized voxels (red). AUC decreases sharply with noise amplitude when the cells are sampled randomly, while it decreases more slowly when the cells are sampled according to phase. The decrease in AUC to chance level (i.e. AUC = 0.5) with the increase in noise amplitude results in insignificant difference in subspace generalization measure (*AUC*_*within-between*_). See supplementary info Figure S3 for the permutation distributions.

How does the difference between the signal variances affect the subspace generalization measure? If the BOLD signal had no noise and all the cells within a voxel were indeed grid cells, the actual variance of the signal would not affect our measure (Figure 2c, the solid and dashed black lines are similar in both panels; i.e. the PCs that explain the activity variance while the agent is in environment one explain the activity variance of environment two similarly well, no matter how the cells are sampled into voxels). However, this is, of course, unrealistic; the BOLD signal is noisy, and it is likely that voxel activity reflects non-grid cells activity as well. To address this, we incorporated noise into our simulated voxel’s activity map. Figure 2c shows that increasing signal variance by grouping according to the grid phase, leads to higher subspace generalization measure (AUC) compared to random sampling; random sampling results in small AUC (*AUC* ≈ 0.5) which is close to the expected AUC following projections on random vectors (solid and dash blue lines in Figure 2c, left, see supplementary info Figure S3 for further analysis). Predictably, as the fraction of randomly sampled grid cells increases the ability to detect subspace generalization in the presence of noise decreases (Figure 2d, Figure S3). Furthermore, sampling of grid cells according to phase increases the statistical power of the subspace generalization method when the amplitude of the noise increases (Figure 2e, Figure S3). To conclude, this shows under noisy conditions, if nearby grid cells have similar phase tuning, as has been shown (Gu *et al*. 2018), our method can in principle detect the generalization properties of grid cells, even in a very low-resolution data, akin to the fMRI BOLD signal. It can in principle work to detect generalization properties of any representation where nearby cells have similar tuning (such as orientation tuning in V1).

### Probing generalisation across abstract tasks with shared statistical rules – task design and behaviour

In human neuroimaging, the success of multivariate pattern analysis (MVPA, (Haxby *et al*. 2001)) and RSA (Kriegeskorte, Mur and Bandettini 2008; Diedrichsen and Kriegeskorte 2017)) tells us that, as with cells, the covariance between fMRI voxel activity contains information about the external world. It is therefore conceivable that we can measure the generalisation of fMRI patterns across related tasks using the same measure of subspace generalisation, but now applied to voxels rather than to cells. This will give us a measure of generlisation in humans that can be used across tasks with no state-to-state mapping – e.g. when the size of the state space is different across tasks. In this section, we first describe the experimental paradigm we used to test whether, as in physical space, EC 1) generalises over abstract tasks governed by the same statistical rules; and 2) does so in a manner that is flexible to the size of the environment. In the next section we use known properties of visual encoding as a sanity check for the use of subspace generalisation on fMRI data in this task. Finally, we describe how the fMRI subspace generalisation results in EC depend on the statistical rules (structural forms) of tasks.

We designed an associative-learning task (Figure 3A and 3B, similar to the task in (Mark *et al*. 2020)) where participants learned pairwise associations between images. The images can be thought of as nodes in a graph (unseen by participants), where the existence of an edge between nodes translates to an association between their corresponding images (Figure 3A). There were two kinds of statistical regularities governing graph structures: a hexagonal/triangular structural form and a community structure. There were also two mutually exclusive image sets that could be used as nodes for a graph, meaning that each structural form had two different graphs with different image sets, resulting in a total of four graphs per participant. Importantly, two graphs of the same structural form were also of different sizes (36 and 42 nodes for the hexagonal structure; 35 and 42 nodes for the community structure - 5 or 6 communities of 7 nodes per community, respectively), meaning states could not be aligned even between graphs of the same structural form. The pairs of graphs with the (approximately) same sizes across structural forms used the same visual stimuli set (Figure 3B). This design allowed us to test for subspace generalisation between tasks with the same underlying statistical regularities, controlling for the tasks’ stimuli and size.

**Figure 3.**
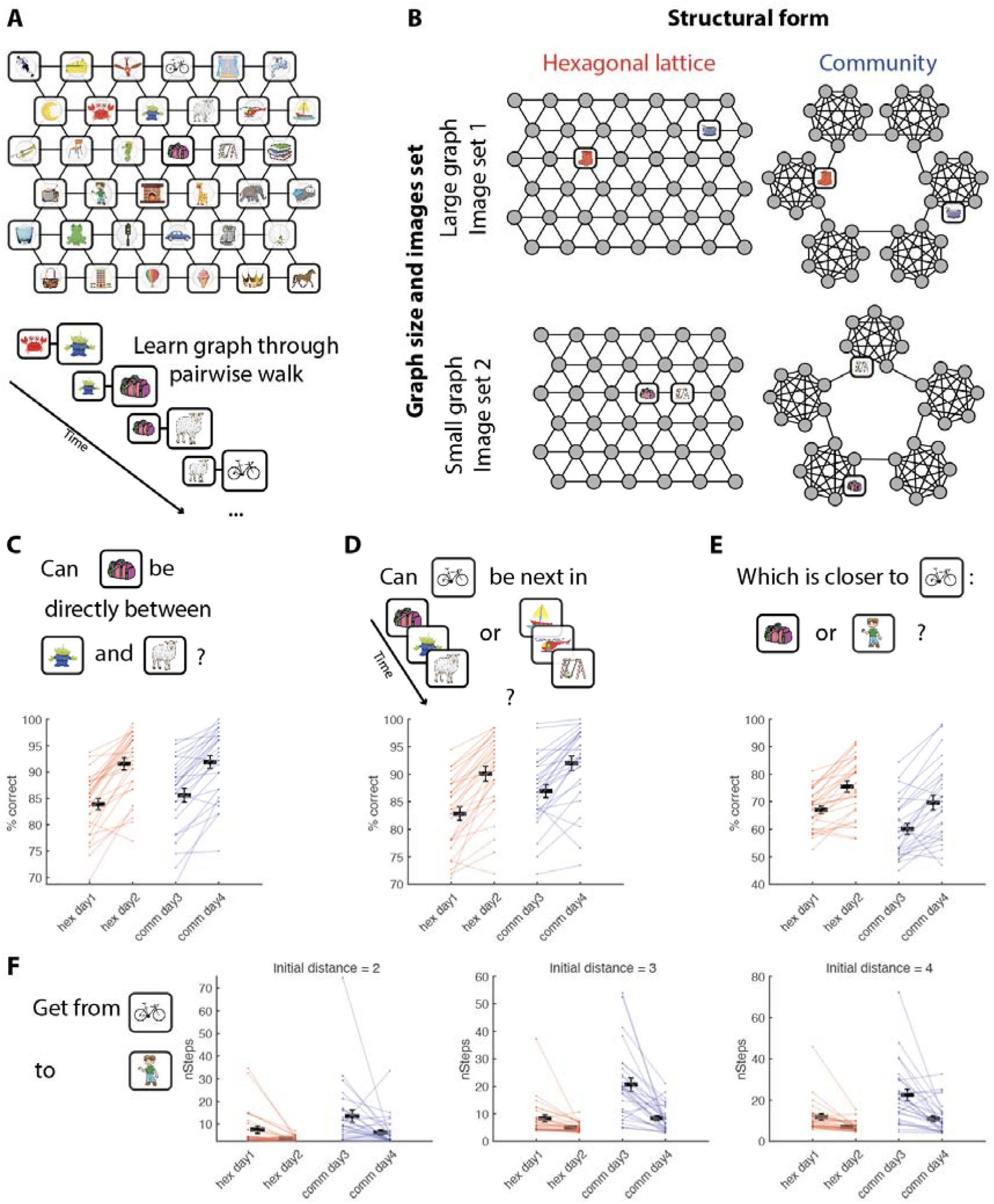
Experimental design and behavior. **A**. Example of an associative graph. Participants were never exposed to this top-down view of the graph - they learned the graph by viewing a series of pairs of neighboring images, corresponding to a walk on the graph. To aid memorisation, we asked participants to internally invent stories that connect the images. **B**. Each participant learned 4 graphs: two with a hexagonal lattice structure (both learned on days 1 and 2) and two with a community structure (both learned on days 3 and 4). For each structural form, there was one larger graph and one smaller graph. The nodes of graphs with approximately the same size were drawn from the same set of images. **C-F**. In each day of training we used four tests to probe the knowledge of the graphs, as well as to promote further learning. In all tests, participants performed above chance level on all days and improved their performance between the first and second days of learning a graph. **C**. Participants were asked whether an image X can appear between images Y and Z (one sided t-test against chance level (50%): hex day1 t(27) = 31.2, p < 10^-22; hex day2 t(27) = 35.5, p < 10^-23; comm day3 t(27) = 26.9, p < 10^-20; comm day4 t(27) = 34.2, p < 10^-23; paired one sided t-test between first and second day for each structural form: hex t(27) = 4.78, p < 10^-5; comm t(27) = 3.49, p < 10^-3). **D**. Participants were shown two 3-long image sequences, and were asked whether a target image can be the fourth image in the first, second or both of the sequences (one sided t-test against chance level (33.33%): hex day1 t(27) = 39.9, p < 10^-25; hex day2 t(27) = 42.3, p < 10^-25; comm day3 t(27) = 44.8, p < 10^-26; comm day4 t(27) = 44.2, p < 10^-26; paired one sided t-test between first and second day for each structural form: hex t(27) = 3.97, p < 10^-3; comm t(27) = 2.81, p < 10^-2). **E**. Participants were asked whether an image X is closer to image Y or image Z, Y and Z are not neighbors of X on the graph (one sided t-test against chance level (50%): hex day1 t(27) = 12.6, p < 10^-12; hex day2 t(27) = 12.5, p < 10^-12; comm day3 t(27) = 5.06, p < 10^-4; comm day4 t(27) = 7.42, p < 10^-07; paired one sided t-test between first and second day for each structural form: hex t(27) = 3.44, p < 10^-3; comm t(27) = 2.88, p < 10^-2). **F**. Participants were asked to navigate from a start image X to a target image Y. In each step, the participant had to choose between two (randomly selected) neighbors of the current image. The participant repeatedly made these choices until they arrived at the target image (paired one sided t-test between number of steps taken to reach the target in first and second day for each structural form. Left: trials with initial distance of 2 edges between start and target images: hex t(27) = 2.57, p < 10^-2; comm t(27) = 2.41, p < 10^-2; MIddle: initial distance of 3 edges: hex t(27) = 2.58, p < 10^-2; comm t(27) = 4.67, p < 10^-2; Right: trials with initial distance of 4 edges: hex t(27) = 3.02, p < 10^-2; comm t(27) = 3.69, p < 10^-3). Note that while feedback was given for the local tests in panels C and D, no feedback was given for the tests in panels E-F to ensure that participants were not directly exposed to any non-local relations. The location of different options on the screen was randomised for all tests. Hex: hexagonal lattice graphs. Comm: community structure graphs.

Participants were trained on the graphs for four days and graph knowledge was assessed in each of the days using a battery of tests described previously (Mark *et al*. 2020 and methods). Some tests probed knowledge of pairwise (neighboring) associations (Figure 3C-D) and others probed “a sense of direction” in the graph, beyond the learned pairwise associations of neighboring nodes (Figure 3 E-F). In all tests, the performance of participants improved with learning and was significantly better than chance by the end of training (Figure 3 C-F), suggesting that participants were able to learn the graphs and developed a sense of direction even though they were never exposed to the graphs beyond pairwise neighbors. Note that while all participants performed well on tests of neighboring associations, the variance across participants for tests of non-neighboring nodes was high, with some participants performing almost perfectly and others close to chance (compare panels C-D to panels E-F). At the end of the training days, we asked participants whether they noticed how the images are associated with each other, 26 out of 28 participants recognized that in two sets, the pictures were grouped.

### FMRI task and analysis

On the fifth day participants performed a task in the fMRI scanner. Each block of the scan included one of the four graphs the participant has learned and started with a self-paced image-by-image random walk on the graph to allow inference of the currently relevant graph (Figure 4a, data not used in this manuscript). The second part of the block had two crucial differences. First, images were arranged into sequences of 3 images that were presented in rapid succession, corresponding to a walk of length 3 on the graph (Figure 4b and Figure S5 for the partitioning the graphs into 3 images sequences). The time between two successive sequences was 800ms (Figure 4c). Second, while the order within each 3-images sequence was dictated by the graph, the order across the sequences was pseudo-random. We needed this second manipulation to ensure coverage of the graph in every block and to eliminate the possibility of spurious temporal correlations between neighboring sequences. However, if we had presented images individually in this random order, graphs with the same stimuli set would have been identical, making it difficult for subjects to maintain a representation of the current graph across the block. Whilst the images were the same across 2 graphs, the sequences of neighboring images uniquely identified each graph, inducing a sensation of “moving” through the graph. To encourage attention to the neighborhood of the sequence in the graph, in 12.5% of trials the sequence was followed by a single image (“catch trial” in Figure 4c), and participants had to indicate whether it was associated with the last image in the sequence (Figure 4c). Participants answered these questions significantly better than chance (Figure S6), indicating that they indeed recognize the correct graph and maintain the correct representation during the block (t-test, p<<0.001 for both structures, t[27]_hex_=11.3, t[27]_comm_=10.6). At the end of each block participants were asked whether they recognised which images set they currently observed (see Method and supplementary for more details). Participants answered these questions significantly better than chance (t-test, p<0.001 for both structures, t[27]_hex_ = 3.8, t[27]_comm_ = 9.96, see supplementary Figure S6), again indicating that they correctly recognised the current graph in the scanner.

**Figure 4.**
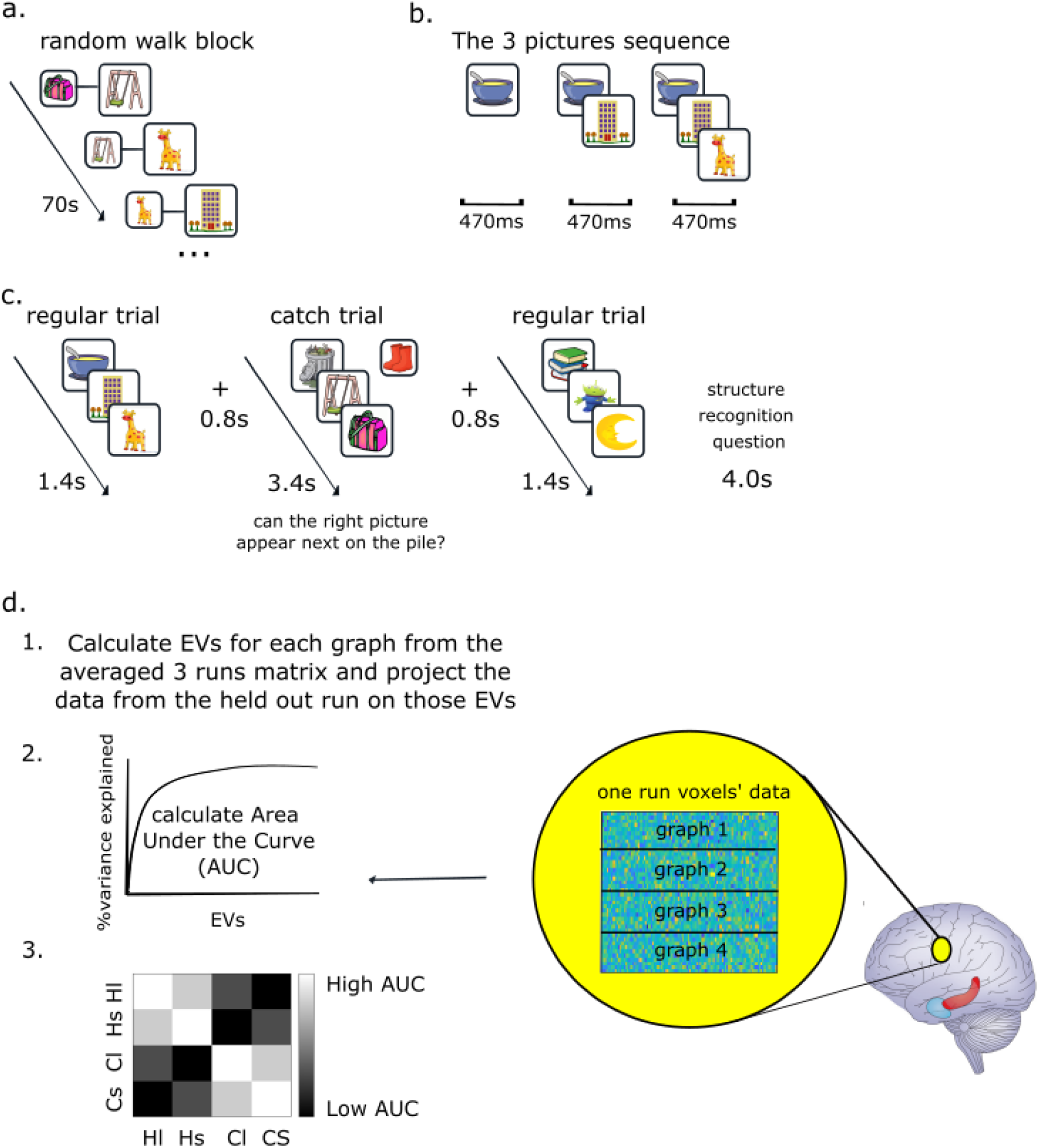
fMRI experiment and analysis method (subspace generalisation) a. Each fMRI block starts with 70s of random walk on the graph: a pair of pictures appears on the screen, each time a participant presses enter a new picture appears on the screen and the previous picture appears behind (similar to the three pictures sequence, sell below). During this phase participants are instructed to infer which “pictures set” (i.e graph) they are currently playing with. Note that fMRI data from this phase of the task is not included in the current manuscript. b. The three pictures sequence: three pictures appear one after the other, while previous picture/s still appear on the screen. c. Each block starts with the random walk (panel a). Following the random walk, sequences of three pictures appear on the screen. Every few sequences there was a catch trial in which we ask participants to determine whether the questioned picture can appear next on the sequence. d. Subspace generalisation method on fMRI voxels. Each searchlight extracts a beta X voxels’ coefficients (of 3-images sequences) matrix for each graph in each run (therefore, there are four such matrices). Then, using cross-validation across runs, the left out run matrix of one graph is projected on the EVs from the (average of 3 runs of the) other graph. Following the projections, we calculate the cumulative percentage of variance explained and the area under this curve for each pair of graphs. This leads to a 4 × 4 subspace generalization matrix that is then being averaged over the four runs (see main text and methods for more details). The colors of this matrix indicate our original hypothesis for the study: that in EC, graphs with the same structure would have larger (brighter) AUCs than graphs with different structures (darker).

To analyze this data, we used the subspace generalisation method as described for the rodent data but replacing the firing of neurons at different spatial locations with the activity of fMRI voxels for different 3-images sequences. To do this, we first performed a voxelwise GLM where each regressor modeled all appearances of a particular 3-images sequence in a given run, together with several nuisance regressors (see Methods). This gave us the activity of each voxel for each sequence. For each voxel, in each run, we extracted the 100 nearest voxels and formed a matrix of sequence X voxels. These are analogous to the data matrices, **B**, in equation 1. We then computed subspace generalisation using the PCs of the voxel X voxel covariance matrix instead of the cell X cell covariance matrix (Figure 4d).

We then employed a leave-one-out cross-validation by repeatedly averaging the activation matrices from three runs of graph X, calculating the PCs from this average representation, and then projecting the activation matrix of the held out run of control graph X (or a test graph Y) on these PCs. This ensures that the “PC” and “data” graphs are always from different runs. We then calculated the subspace generalisation between each pair of graphs resulting in a 4×4 matrix at each voxel of the brain (Figure 4d).

We refer to the elements of this 4×4 matrix in the following notation: we denote by H/C graphs of either hexagonal or community structure, and by s/l either small or large stimuli sets (matched across graphs of different structures). For example, HsCs denotes the element of the matrix corresponding to activity from the small hexagonal graph projected on PCs calculated from the small (same image-set) community-structure graph.

### Testing subspace generalisation on visual representations

To verify our analysis approach is indeed valid when used on our fMRI data, we first tested it on the heavily studied object encoding representations in lateral occipital cortex (LOC, Malach 1995 PNAS, Grill-Spector). Recall that our stimuli in the scanner were concurrently presented sequences of three images of objects. We reasoned that these repeated sequences would induce correlations between object representations that should be observable in the fMRI data and detectable by our method. This would allow us to identify visual representations of the objects without ever specifying when the stimuli (i.e. 3-images sequences) were presented.

To this end we compared subspace generalization computed between different runs that included the same stimuli (3-images sequences, with different order across sequences between runs) with subspace generalization computed between runs of different stimuli while controlling for the graph structure. This led to the contrast [HlHl + ClCl + HsHs +CsCs] - [HlHs + HsHl + ClCs + CsCl], which had a significant effect in LOC (Figure 5a, peak MNI [−44,−86,−8], t(27)_peak = 4.96, P_tfce < 0.05 based on a FWE-corrected nonparametric permutation test, corrected in bilateral LOC mask (Harvard-Oxford atlas, Desikan 2006, Neuroimage). In an additional exploratory analysis, we tested the significance of the same contrast in a whole-brain searchlight. While this analysis did not reach significance once corrected for multiple comparisons, the strongest effect was found in LOC (Figure 5a). Note that in this contrast we intentionally ignored the elements of the 4×4 matrix where the data and the PCs came from graphs with the same images set and a different structure (HlCl, HsCs, ClHl, CsHs), because they did not share the exact same visual stimuli (the 3-images sequence). In these cases, we did not have a hypothesis about the subspace generalization in LOC. These results suggest that we can detect the correlation structure induced by stimuli without specifying when each stimulus was presented.

**Figure 5:**
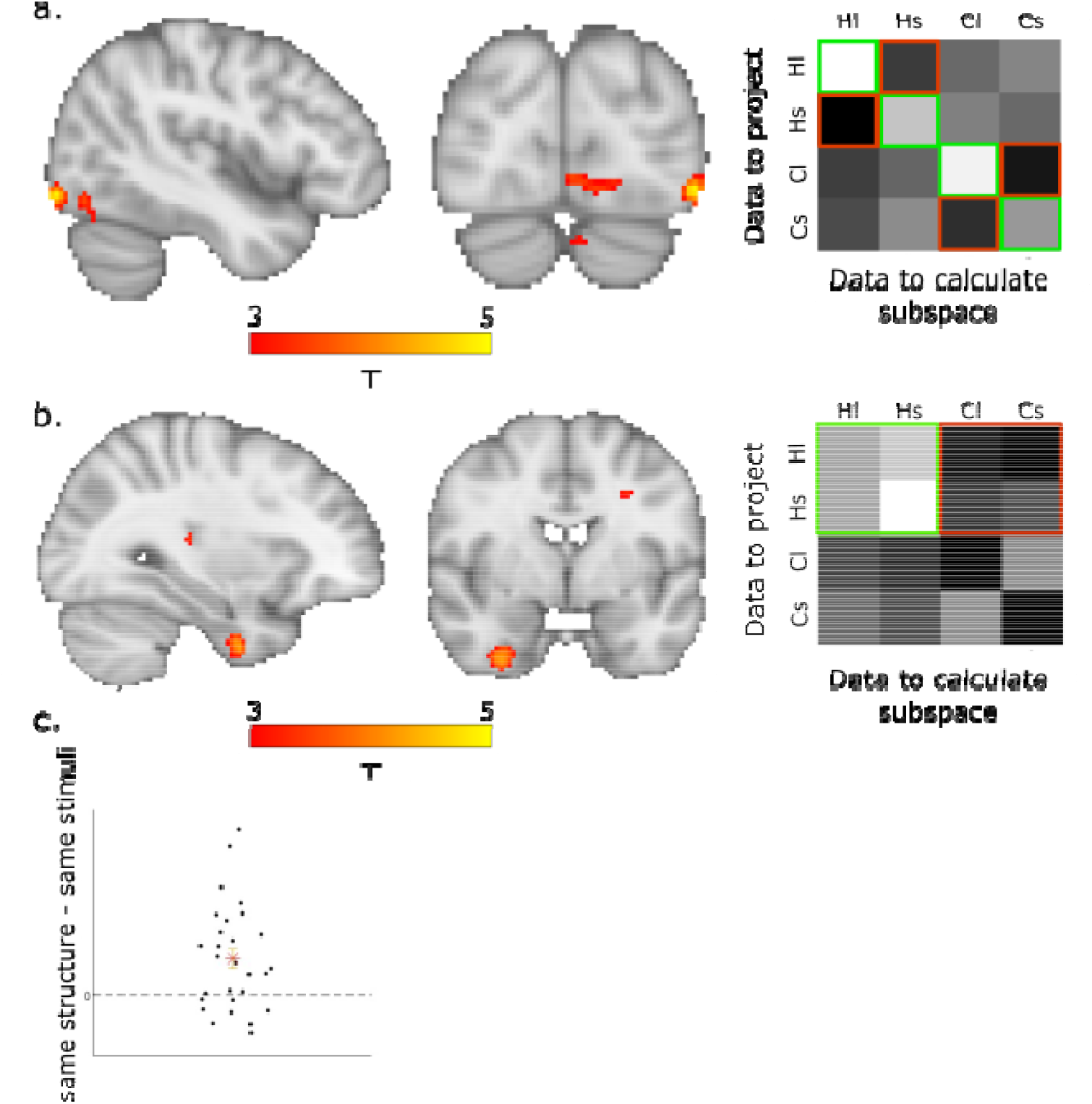
subspace generalisation in visual and structural representations. a. Subspace generalisation of visual representations in LOC. Left: difference in subspace generalization was computed between different blocks that included the same stimuli with subspace generalization computed between blocks of different stimuli while controlling for the graph structure, i.e [HlHl + ClCl + HsHs +CsCs] - [HlHs + HsHl + ClCs + CsCl]. t(27)_peak = 4.96, P_tfce < 0.05 over LOC. Right: visualization of the subspace generalisation matrix (averaged over all LOC voxels with t>2 for the [HlHl + ClCl + HsHs +CsCs] - [HlHs + HsHl + ClCs+ CsCl] contrast, i.e. green minus red entries. b. EC generalises over the structure of hexagonal graphs. Left: the effect for the contrast [HlHl + HlHs + HsHl + HsHs] - [HlCl + HlCs + HsCl + HsCs], i.e. the difference between subspace generalisation of hexagonal graphs data, when projected on PCs calculated from (cross-validated) hexagonal graphs (green elements in right panel) vs community structure graphs (red elements). t(27)_peak = 4.2, P_tfce <0.01 over EC. Right: Same as in a. right but for the [HlHl + HlHs + HsHl + HsHs] - [HlCl + HlCs + HsCl + HsCs] contrast in EC. c. The average effect in an ROI from Baram et al. (green cluster in figure 3d of Baram et al.) for each participant. Star denotes the mean, error bars are SEM.

### EC generalizes a low-dimensional representation across hexagonal graphs of different stimuli and sizes

Having established that the subspace generalization method can detect meaningful correlations between fMRI voxels, we next aimed to test whether EC will represent the statistical structure of abstract graphs with generalisable low-dimensional representations. We first tested this for discretized 2D (hexagonal) graphs, using the community structure graphs as controls: We tested whether the EC subspaces from hexagonal graphs blocks were better aligned with the PCs of other hexagonal blocks, than with the PCs from community graphs blocks, i.e. ([HlHl + HlHs + HsHl + HsHs] - [HlCl + HlCs + HsCl + HsCs], Figure 5b). This contrast was significant in the right EC (peak MNI [28, −10, −40], t(27)_peak = 4.2, P_tfce <0.01 based on a FWE-corrected nonparametric permutation test, corrected in a bilateral EC mask (Figure 5b) (Julich atlas, Eickoff 2007). We obtained a null result for the equivalent analysis for community structure graphs ([ClCl + ClCs + CsCl + CsCs] - [ClHl + ClHs + CsHl + CsHs]). This was particularly due to low subspace generalization across different runs of the same community structure graphs (bottom two diagonal elements in Figure 5b right, compare to our original hypothesis subspace generalization matrix in Figure 4d). See the Discussion for possible interpretations of this null result.

To ensure the robustness of the hexagonal graphs result we next tested the same effect in an orthogonal ROI from our previous study. In (Baram *et al*. 2021) we have shown that EC generalises over different reinforcement learning tasks with the exact same structure. We therefore tested the same effect in that ROI (all voxels in the green cluster in Figure 3d in Baram 2021 *et al*., peak MNI: [25, −5, −28]), and indeed the [HlHl + HlHs + HsHl + HsHs] - [HlCl + HlCs + HsCl + HsCs] contrast was significant (one sided t-test, t(27) =3.6, p<0.001, Figure 5c).

Taken together, these results suggest that as in physical space, different abstract hexagonal graphs are being represented on the same EC low-dimensional subspace. This is consistent with a view where the same EC cell assembly represents both hexagonal graphs, and that these cells covary together - even when the underlying size of the graph is different.

## Discussion

The contributions of this manuscript are two-fold: first, we show that EC representations generalize over hexagonal abstract graphs of different sizes, highlighting the importance of the statistical properties of the environment to generalization. This expands our previous work (both experimental (Baram *et al*. 2021) and theoretical (Whittington *et al*. 2020)), suggesting EC plays an important role in generalization over abstract tasks, to the case where the tasks are governed by the same statistical rules but are not governed by the exact underlying graph (transition structure). This view builds on the known generalization properties of EC in physical space (Fyhn *et al*. 2007; Gardner *et al*. 2022) and on recent literature highlighting parallels between medial temporal lobe representations in spatial and non-spatial environments (Behrens *et al*. 2018; Whittington *et al*. 2022). Second, we present an fMRI analysis method (“subspace generalization”), adapted from related work in electrophysiology analysis (Samborska *et al*. 2022), to quantify generalization in cases where a mapping between states across environments is not available (though see (Hahamy and Behrens 2019) for our previous fMRI application of this method in the visual domain).

Exploiting previous knowledge while making decisions in new environments is a hard challenge that humans and animals face regularly. To enable generalization from loosely related previous experiences, knowledge should be represented in an abstract and flexible manner that does not depend on the particularities of the current task. Understanding the brain’s solution to this computational problem requires a definition of a “generalisable representation”, and a way of quantifying it. Here, we define generalization as sharing of neuronal manifold across representations of related tasks. The particular assumption here is that in the EC, such manifolds encode the relevant information about the particular structural form of the task.

An example of such generalization has previously been observed in the spatial domain, in grid cells recordings across different physical environments, regardless of shape or size (Fyhn *et al*. 2007; Gardner *et al*. 2022). This was usually done through direct comparison of the pairwise activity patterns of cells (Fyhn *et al*. 2007; Yoon *et al*. 2013; Gardner *et al*. 2022). However, this is not possible to do in fMRI, rendering the examination of EC generalization in complex abstract tasks difficult. “Subspace generalization” relies on the idea that similarity in activity patterns across tasks implies similarity of the within-task correlations between neurons. These are summarized in the similarity between the (low dimensional) linear subspaces where the activity of the neurons/voxels representing the two tasks lies. For fMRI purposes, this similarity between within-task neuronal correlations should be reflected in the similarity between within-task correlations across voxels, as long as the relevant neurons anatomically reside across a large enough number of voxels. Importantly, comparing similarity in neuronal correlations structures rather than similarity in states representations patterns (as in RSA) allows us to examine flexible knowledge representations when a mapping between states in the two tasks does not exist. We present three validations of this method: in cells, we show it captures all expected properties of grid and place cells, even if we reduce the data resolution by averaging over the activity of group of cells. In simulation, we show that calculating subspace generalization using simulated voxels from simulated grid cells results in significant generalization effect under realistic condition. In fMRI, we show it captures the expected correlations induced by the visual properties of a task in LOC.

Our main finding of subspace generalization in EC across hexagonal graphs with different sizes and stimuli significantly strengthens the suggestion that EC flexibly represents all ‘spatial-like’ tasks, such as discretized 2D hexagonal graphs. Recently, we presented a theoretical framework for this idea: a neural network trained to predict future states, that when trained on 2D graphs displayed known spatial EC representations (the Tolman Eichenbaum Machine (TEM) (Whittington *et al*. 2020)). However, ‘spatial-like’ structures are not the only prevalent structures in natural tasks. The relations between task states often follow other structural forms (such as periodicities, hierarchies or community structures), inference of which can aid behavior (Mark *et al*. 2020). Representations of non-Euclidian task structures have been found in EC (Garvert, Dolan and Behrens 2017; Baram *et al*. 2021) and these generalize over different reinforcement learning tasks that are exactly the same except for their sensory properties (Baram *et al*. 2021). Indeed, when TEM was trained on non-Euclidean structures like hierarchical trees, it learned representations that were generalisable to novel environments with the same structure (Whittington *et al*. 2020). Further, we have previously shown that representing each family of graphs of the same structural form with the relevant stable representation (i.e. basis set) allows flexible transfer of the graph structure and therefore inference of unobserved transitions (relations between task’s states) (Mark *et al*. 2020). Together these studies suggest that flexible representation of structural knowledge may be encoded in the EC.

Based on these, we hypothesized that EC representations will also generalize over non-‘spatial-like’ tasks (here, community-structure) of different sizes. However, we could not find conclusive evidence for such a representation: the relevant contrast ([ClCl + ClCs + CsCl + CsCs] - [ClHl + ClHs + CsHl + CsHs]) did not yield a statistically significant effect in EC (or elsewhere, in an exploratory analysis corrected across the whole brain). This is despite clear behavioral evidence that participants use the community structure of the graph to inform their behavior: participants have a strong tendency to choose to move to the connecting nodes (nodes that connect two different communities) over non-connecting nodes ((Mark *et al*. 2020), and Figure S4a). Moreover, in the post-experiment debriefing, participants could verbally describe the community structure of the graphs (26 out of 28 participants). This was not true for the hexagonal graphs. Why, then, did we not detect any neural generalization signals for the community structure graphs? There are both technical and psychological differences between the community structure and the hexagonal graphs that might have contributed to the difference in the results between the two structures. First, we have chosen a particular nested structure in which communities are organized on a ring. Subspace generalisation may not be suitable for the detection of community structure: for example, a useful generalisable representation of such structure is composed of a binary ‘within-community nodes’ vs ‘connecting nodes’ representation. If this is the representation used by the brain, it means all “community-encoding” voxels are similarly active in response to all stimuli (as all 3-images sequences contain at least two non-connecting node images), and only “connecting nodes encoding” voxels change their activation during stimuli presentation. Therefore, there is very little variance to detect.

Though this manuscript has focused on EC, it is worth noting that there is evidence for structural representations in other brain areas. Perhaps the most prominent of these is mPFC, where structural representations have been found in many contexts (Klein-Flügge *et al*. 2019; Baram *et al*. 2021; Klein-Flügge, Bongioanni and Rushworth 2022). Indeed, the strongest grid-like signals in abstract 2D tasks are often found in mPFC (Constantinescu, O’Reilly and Behrens 2016; Bao *et al*. 2019; Park *et al*. 2020; Bongioanni *et al*. 2021) and task structure representations have been suggested to reside in mOFC (Wilson *et al*. 2014; Schuck *et al*. 2016; Xie and Padoa-Schioppa 2016). The difference and interaction between PFC and MTL representations is a very active topic of research. One such suggested dissociation that might be of relevance here is the preferential contribution of MTL and PFC to latent and explicit learning, respectively. A related way of discussing this dissociation is to think of mPFC signals as closer to the deliberate actions subjects are taking. Circumstantial evidence from previous studies in our lab (tentatively) suggest the existence of such dissociation also for structural representations: when participants learnt a graph structure without any awareness of it, this structure was represented in MTL but not mPFC (Garvert, Dolan and Behrens 2017). On the other hand, when participants had to navigate on a 2D abstract graph to locations they were able to articulate, we observed much stronger grid-like signals in mPFC than MTL (though a signal in EC was also observed,(Constantinescu, O’Reilly and Behrens 2016)). In addition, Baram *et al*. found that while the abstract structure of a reinforcement learning task was represented in EC, the structure-informed learning signals that inform trial-by-trial behavior with generalisable information were found in mPFC. Taken together, these results suggest that here, it is reasonable to expect generalisation signals of community structure graphs (of which participants were aware) in PFC, as well as the signals reported in EC for hexagonal graphs (of which participants were unaware). Indeed, when we tested for subspace generalisation of community structure graphs in the same ROI in vmPFC where Baram *et al*. found generalisable learning signals, we obtained a significant result (though this is a weak effect, and we hence report it with caution in the supplementary material, Figure S4b).

To summarize, we have extended the understanding of EC representations and showed that EC represents hexagonal graph structures of different sizes, similarly to grid cells representation of spatial environments. We did this by using an analysis method which we believe will prove useful for the study of generalisable representations in different neural recording modalities. More work is needed to verify whether this principle of EC representations extends to other, non-’’spatial-like’ structural forms.

## Methods

### Rodent cells analysis

Cells electrophysiology data were taken from (Chen *et al*. 2018). In short, cells (place cells from CA1 and grid cells from dmEC) were recorded while the animals foraged in two different square arenas; one real arena and one virtual reality (VR) arena, real arena is 60×60 and the VR arena is 60×60 or 90×90 cm. The VR system restrained head-movements to horizontal rotations, and included an air-suspended ball on which the mice could run and turn. A virtual environment reflecting the mouse’s movements on the ball was projected on screens in all horizontal directions and on the floor. Mice were implanted with custom-made microdrives (Axona, UK), loaded with 17mm platinum-iridium tetrodes, and providing buffer amplification. We analyzed grid cells data from three animals; two animals had only grid cells data and one animal had both place cells and grid cells data. We analyzed place cells data from three more animals that had only place cells data (mouse 1 had 14 grid cells, mouse 2 and 3 had 21 grid cells, mouse 1, 4, 5 had 25 place cells). This experimental design results in two different firing rate maps, one for each arena. After preprocessing (calculate the firing rate map using on 64×64 bins matrix and smoothing of the firing rate maps with 5 bins boxcar), we calculated the ‘subspace generalisation’ score, as follows:

a. Calculate the neuron X neuron correlation matrix from the first firing rate map (one of the environments) and its principal components (PCs).
b. Project the firing rate maps from this environment and the other environment on these PCs.
c. Calculate the cumulative variance explained as a function of PCs (that are organized according to their corresponding eigenvalues)
d. Calculate the area under the curve (AUC).

*Permutation test* 1 (within cell type): Our hypothesis is that the neuron X neuron correlation structure is preserved while the animals forage in the two different arenas, i.e. that the active cells’ assemblies remain the same. Therefore, the null hypothesis is that the cells’ assemblies are random and did not remain the same while animals forage in the two arenas. We therefore calculated the PCs using the firing rate map while the animal foraged in one environment and permuted the cells’ identity of the firing rate maps correspond to the second environment. We then calculated the difference between the ‘subspace generalisation’ score within and across environments. This creates our null distribution, which we compare to the subspace generalisation score of the non-permuted data.

*permutation test* 2 (between cell types): Our hypothesis is that grid cells generalise better than place cells, i.e. that the difference between the AUC of within arena projection to across arenas projection is smaller in grid cells compared to place cells. To this end, we created AUC-differences distribution using place cells activity as our null distribution; we sample place cells from each animal, such that the number of grid cells and place cells was equal (mouse 1 had 14 grid cells, mouse 2 and 3 had 21 grid cells, mouse 1, 4, 5 had 25 place cells). Then, for each sample, we calculated the difference in AUC (same arena - different arenas), as before. We calculated the distribution of these AUC-differences values from all three animals. We then checked whether the AUC-differences in grid cells, for all three animals, is significantly smaller than those predicted by the sampled place cells distribution (Figure S1).

#### Reducing the resolution of the electrophysiological data

We first normalized all firing rate maps. Then, for each animal we randomly sampled (with repeats) seven cells into two groups and averaged the cells’ activity within each group, separately for each environment. We then concatenated the resulted size-2 vectors from all animals into one vector and used this vector as above to calculate the AUC differences between within and across environments. The number of bootstraps was 400, therefore we had 800 repetitions to calculate the distribution (for each sample we project on both environments therefore getting two AUC - difference values). The plots in Figure.1d were smoothed with smoothing window of 9, the number of bins to calculate the distribution was 50.

#### Simulating pseudo voxels

Grid cells are simulated as a thresholded sum of three 2D cosines (Burgess *et al*. 2007). Each module is simulated by shifting the grid cells within a grid that spans the rhombus of the hexagonal grid, such that the average over all grid cells within a module is a constant across the box (note that due to numerical issues this is almost constant).

We simulated 13456 cells per module (116*116 in the x-y plane, i.e. covering the grid’s rhombus). The box is simulated with 50*50 resolution (the size of the “box” is 10*10). We simulated four different modules that differ in their grid spacing and phases. Each environment was simulated by a different phase and shift of the grid fields such that the relationships between the cells remain the same across environments.

Voxels were simulated by averaging cells within a module. Each module was segregated into four groups of cells (therefore there are 3364 cells within each voxel, see supplementary for different segregations). Each voxel is an average over the cells’ firing rate map within the group. The averaging was done in two stages:

a. sampling grid cells randomly - i.e. not related to their grid phase

b. The remaining cells were segregated into four groups according to their phase.

The above process was repeated for different fractions of random/(according to phase) ratio (*ratio_random* = [0,1], 0: only segregated according to phase, 1: only segregated randomly). We further added spatial white noise to each voxel, noise std ranging from 0 to 0.1. When examining the effect of random sampling, the noise std was 0 or 0.1.

### FMRI experiment

#### Participants

60 UCL students were originally recruited. As the training is long and hard, for each scan we recruited two participants for the training sessions, and chose the better performing of the two to be scanned. Overall, we scanned 34 participants and excluded 6 participants from the analysis because of severe movement or sleepiness in the scanner.

The study was approved by the University College London Research Ethics Committee (Project ID 11235/001). Participants gave written informed consent before the experiment.

### Behavioural training for fMRI training task

To ensure that participants understood the instructions, the first training day was performed in the lab while the other three training days were performed from the participant’s home.

### Graphs

One hexagonal graph consisted of 36 nodes and the other 42 nodes as shown in Figure 3b. One community structured graph consisted of 5 communities and the other 6 communities, with 7 nodes each. Within a community, each node was connected to all other nodes except for the two connecting nodes that were not connected to each other but were each connected to a connecting node of a neighboring community (Figure 3b). Therefore, all nodes had a degree of six, similarly to the hexagonal graphs (except the nodes on the hexagonal graphs border, which had degree less than six). Our community structure graph had a hierarchical structure, wherein communities were organized on a ring.

#### Training procedures

In each of the training days, participants learned two graphs with the same underlying structure but different stimuli. During the first two days participants learned the hexagonal graphs, while during the third and fourth days participants learned the community structured graphs. We chose to first teach the hexagonal graphs structure for all participants and not randomize the order because learning community structure graph changes participants’ learning strategy (mark *et al*. 2020). During the fifth day, before the fMRI scan, participants were reminded of all four graphs, with two repetitions of each hexagonal graph and one repetition of each community structured graph. Stimuli were selected randomly, for each participant, from a bank of stimuli (each pair of graphs, one hexagonal and one of a community structured graph shared the same bank). Each graph was learnt during four blocks (Figure. 3b; 4 blocks for graph 1 followed by 4 blocks for graph 2 in each training day). Participants could take short resting breaks during the blocks. They were instructed to take a longer resting break after completing the four blocks of the first graph of each learning day.

#### Block structure

Each block during training was made of the following tasks: 1) Learning phase 2) Extending pictures sequences 3) Can it be in the middle 4) Navigation 5) Distance estimation (see Figure 3). Next, we elaborate the various components of each block.

#### Learning phase (Figure 3a)

Participants learned associations between graph nodes by observing a sequence of pairs of pictures which were sampled from a random walk on the graph (successive pairs of pictures shared a common picture). Participants were instructed to ‘say something in their head’ in order to remember the associations. Hexagonal graphs included 120 steps of the random walk per block and community-structured graphs included 180 steps per block (we introduced more pictures in the community graph condition as random walks on such graphs result in high sampling of transitions within a certain community and low sampling of transitions between communities).

#### Extending pictures sequences (Figure 3d)

Given a target picture, which of two sequences of three pictures can be extended by that picture (a sequence can be extended by a picture only if it is a neighbor of the last picture in the sequence, the correct answer can be sequence 1/sequence 2/both sequences): Sixteen questions per block. (note that a picture could not appear twice in the same sequence, i.e. if the target picture is already in the sequence the correct answer was necessarily the other sequence).

#### Can it be in the middle (Figure 3c)

Determine whether a picture can appear between two other pictures, the answer is yes if and only if the picture is a neighbor of the two other pictures. Sixteen questions per block.

#### Navigation (Figure 3e)

The aim—navigate to a target picture (appears at the right of the screen). The task was explained as a card game. Participants are informed that they currently have the card of the picture that appears on the left of the screen. They were asked to choose between two pictures that are associated with their current picture. They could also skip and sample again two pictures that are associated with the current picture, if they thought their two current options did not get them closer to the target (skipping was counted as a step). In each step participants were instructed to choose a picture that they thought had a smaller number of steps to the target picture (according to their memory). Following choice, the chosen picture appeared on the left and two new pictures, that correspond to states that are neighbors of the chosen picture, appear as new choices. After a participant selected a neighbor of the target picture, that target picture itself could appear as one of the new options for choice. The game terminated when either the target was reached or 200 steps were taken (without reaching the target). In the latter case a message ‘too many steps’ was displayed. On the first block, for each step, the number of links from the current picture to the target picture was shown on the screen. Participants played three games (i.e. navigation until the target was reached or 200 steps passed) in each block, where the starting distance (number of links) between the starting picture to the target was 2, 3 and 4.

#### Distance estimation

Which of two pictures has the smallest number of steps to a target picture: 45 questions per block (none of the 2 pictures was a direct neighbor on the graph, i.e. the minimal distance was 2 and no feedback was given).

### fMRI scanning task

The task consisted of four runs. Each run was divided into five blocks (one block for each graph and one more repetition for one of the hexagonal graphs; the repetition was not used in the analyses in this manuscript). On each block participants observed pictures that belong to one of the graphs. A block started with 70sec in which participants observed, at their own pace, a random walk on the graph; two neighboring pictures appeared on the screen and when participants pressed ‘enter’ a new picture appeared on the screen (similar to the training learning phase). The new picture appeared in the middle of the screen and the old picture appeared on its left. Participants were instructed to infer which ‘pictures set’ they are currently observing. No information about the graph was given. This random walk phase was not used in any analyses in this manuscript. Next, sequences of three pictures appeared on the screen, one after the other (note the first and second pictures did not disappear from the screen until after the third picture in the sequence was presented - all three pictures disappeared together, prior to the next trial, Figure 4b). To keep participants engaged, once in a while (5 out of 45 sequences) a fourth picture appeared and participants had to indicate whether this picture can appear next on the sequence (‘catch trials’, Figure 4c). Before starting the fMRI scan participants were asked whether they found any differences between the picture sets during the first two days (when the hexagonal graphs were learnt) and the last two days (when the community graphs were learnt). Most participants (26 out of 28) could indicate that there were groups of pictures (i.e. communities) in the last two days, and that this was not the case during the first two days. At the end of each block in the scanner participants answered whether or not there are groups in the current picture set (participants that were not aware of the groups were asked whether this set belongs to the first two training days or not). Participants were given a bonus for answering correctly, such that 100% correct results in a ten pounds bonus.

### fMRI data acquisition

FMRI data was acquired on a 3T Siemens Prisma scanner using a 32 channels head coil. Functional scans were collected using a T2*-weighted echo-planar imaging (EPI) sequence with a multi-band acceleration factor of 4 (TR = 1.450 s, TE = 35ms, flip angle = 70 degrees, voxel resolution of 2×2×2mm). A field map with dual echo-time images (TE1 = 10ms, TE2 = 12.46ms, whole-brain coverage, voxel size 2×2×2mm) was acquired to correct for geometric distortions due to susceptibility-induced field inhomogeneities. Structural scans were acquired using a T1-weighted MPRAGE sequence with 1×1×1mm voxel resolution. We discarded the first six volumes to allow for scanner equilibration.

### Pre-processing

Pre-processing was performed using tools from the fMRI Expert Analysis Tool (FEAT, Woolrich MW *et al*. 2001; Woolrich MW et al. 2004), part of FMRIB’s Software Library (FSL, Smith et al. 2004). Data from each of the four scanner runs was preprocessed separately. Each run was aligned to a reference image using the motion correction tool MCFLIRT. Brain extraction was performed using the automated brain extraction tool BET (Smith, 2002). All data were temporally high-pass filtered with a cut-off of 100s. Registration of EPI images to high-resolution structural images and to standard (MNI) space was performed using FMRIB’s Linear Registration Tool (FLIRT (Jenkinson et al., 2002; Jenkinson and Smith, 2001)). No spatial smoothing was performed during pre-processing (see below for different smoothing protocols for each analysis). Because of the notable breathing- and susceptibility-related artifacts in the entorhinal cortex, we cleaned the data with FMRIB’s ICA tool, FIX (Griffanti et al. 2014; Salimi-Khorshidi *et al*. 2014).

### Univariate analysis

Due to incompatibility of FSL with the MATLAB RSA toolbox (Nili *et al*. 2014) used in subsequent analyses, we estimated all first-level GLMs and univariate group-level analyses using SPM12 (Wellcome Trust Centre for Neuroimaging, https://www.fil.ion.ucl.ac.uk/spm).

For estimating subspace generalization, we constructed a GLM to estimate the activation as a result of each three images’ sequence (a ‘pile’ of pictures). The GLM includes the following regressors: mean CSF regressor and 6 motion parameters as nuisance regressors, bias term modeling the mean activity in each fMRI run, a regressor for the ‘start’ message (as a delta function), a regressor for the self-paced random walk on each graph (a delta function for each new picture that appears on the screen), a regressor for each pile in each graph (duration of a pile: 1.4sec), regressor for the catch trial onset (delta) and the pile that corresponds to the catch (pile duration). All regressors beside the 6 motion regressors and CSF regressor were convolved with the HRF. The GLM was calculated using non-normalized data.

### Multivariate analysis

#### Quantifying subspace generalization

We calculated noise normalized GLM betas within each searchlight using the RSA toolbox. For each searchlight and each graph, we had a nVoxels (100) by nPiles (10) activation matrix (*B*_*voxel*x*pile*_) that describes the activation of a voxel as a result of a particular pile (three pictures’ sequence). We exploited the (voxel x voxel) covariance matrix of this matrix to quantify the manifold alignment within each searchlight.

To account for fMRI auto-correlation we used Leave One Out (LOO) approach; For each fMRI scanner run and graph, we calculated the mean activation matrix over the three others scanner runs 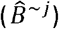. We then calculated the left Principal Component (PCs) of that matrix 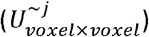. To quantify the alignment, we projected the excluded scanner run graph activation matrix (*B*^*j*^) of each graph on these PCs and calculated the accumulated variance explained as a function of PCs, normalized by the total variance of each graph within each run. Therefore, for each run and graph we calculated:

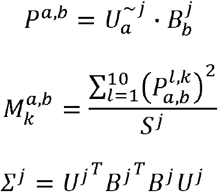

Where *P*^*a,b*^ is the projection matrix of dimensions *voxel* × *pile* of graph *‘b’* on the PCs of graph *‘a’*, 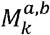 is the normalized variance explained on the *‘k’* direction, *s*^*j*^ is the summation of the diagonal of *∑* ^*j*^, the total variance as a result of the graph piles (three images sequence). We then calculated the cumulative variance explained over all *‘k’* PCs directions. As a summary statistic we calculated the area under this curve. This gives us a 4×4 alignment matrix, for each run, such that each entry (*a, b*) in this matrix is a measure of the alignment of voxels patterns as a result of the two graphs *a&b* (*Figure 4d*). We then averaged over the four runs and calculated different contrasts over this matrix.

The above calculations were performed in subject space, we therefore normalized the searchlight results and then smoothed with a kernel of 6mm FWHM using FSL FLIRT and FNIRT before performing group level statistics.

For group level we calculated the t-stat over participants of each contrast:

Visual contrast was [HlHl + ClCl + HsHs +CsCs] - [HlHs + HsHl + ClCs + CsCl], i.e. same exact sequences controlled by the same structure.

Structural contrast was [HlHl + HlHs + HsHl + HsHs] - [HlCl + HlCs + HsCl + HsCs], i.e. the difference between subspace generalisation of hexagonal graphs data, when projected on PCs calculated from (cross-validated) hexagonal graphs (yellow elements in middle panel) vs community structure graphs (red elements).

#### Multiple comparisons correction

Multiple comparison correction was performed using the permutation tests machinery (Nichols and Holmes 2002) in PALM (Winkler *et al*. 2014): within the mask we used for multiple comparisons correction (details in main text), we first measured the TFCE statistic for the current contrast. We then repeated this procedure for each of the 10000 random sign-flip iterations (each participant’s contrast sign was randomly flipped and the statistic over participants was calculated). Using these values we then created a null distribution of TFCE statistics by saving only the voxel with the highest TFCE in each iteration. Comparing the true TFCE to the resulting null distributions results in FWE-corrected TFCE P-values.

The code for the analysis and simulation is in: https://github.com/ShirleyMgit/subspace_generalization_paper_code/tree/main

## Author contributions

S.M and T.B conceive the research, S.M and P.S design and perform the experiment, S.M and A.B analyzed the data. A.H and V.S provided consultation with analysis. S.M, A.B and T.B wrote the manuscript.

## Acknowledgements

T.B. is supported by a Wellcome Principal Research Fellowship (219525/Z/19/Z), a Wellcome Collaborator award (214314/Z/18/Z), for the purpose of Open Access, the author has applied a CC BY public copyright licence to any Author Accepted Manuscript version arising from this submission, a JS McDonnell Foundation award (JSMF220020372), and by the Jean Francois and Marie-Laure de Clermont Tonerre Foundation. The Wellcome Centre for Integrative Neuroimaging and Wellcome Centre for Human Neuroimaging are each supported by core funding from the Wellcome Trust (203139/Z/16/Z, 203147/Z/16/Z). The Sainsbury-Wellcome centre is supported by core funding from the Wellcome Trust (219627/Z/19/Z) and the Gatsby Charitable Foundation (GAT3755).

A.H. was supported by the European Molecular Biology Organization nonstipendiary Long-Term Fellowship (848–2017), Human Frontier Science Program (LT000444/2018), Israeli National Postdoctoral Award Program for Advancing Women in Science, and the European Union’s Horizon 2020 research and innovation programme under the Marie Skłodowska-Curie Grant Agreement No. 789040.

## References

1. Bao X, Gjorgieva E, Shanahan LK et al. Grid-like Neural Representations Support Olfactory Navigation of a Two-Dimensional Odor Space. Neuron 2019;102:1066–1075.e5.

2. Baram AB, Muller TH, Nili H et al. Entorhinal and ventromedial prefrontal cortices abstract and generalize the structure of reinforcement learning problems. Neuron 2021;109:713–23.

3. Barron HC, Garvert MM, Behrens TE. Repetition suppression: a means to index neural representations using BOLD? Philos Trans R Soc B Biol Sci 2016;371:20150355.

4. Behrens TE, Muller TH, Whittington JC et al. What is a cognitive map? Organizing knowledge for flexible behavior. Neuron 2018;100:490–509.

5. Bongioanni A, Folloni D, Verhagen L et al. Activation and disruption of a neural mechanism for novel choice in monkeys. Nature 2021;591:270–4.

6. Burak Y, Fiete IR. Accurate path integration in continuous attractor network models of grid cells. PLoS Comput Biol 2009;5:e1000291.

7. Burgess N, Barry C, O’keefe J. An oscillatory interference model of grid cell firing. Hippocampus 2007; 801–812, 17(9).

8. Chen G, King JA, Lu Y et al. Spatial cell firing during virtual navigation of open arenas by head-restrained mice. Elife 2018;7:e34789.

9. Constantinescu AO, O’Reilly JX, Behrens TE. Organizing conceptual knowledge in humans with a gridlike code. Science 2016;352:1464–8.

10. Diedrichsen J, Kriegeskorte N. Representational models: A common framework for understanding encoding, pattern-component, and representational-similarity analysis. PLOS Comput Biol 2017;13:e1005508.

11. Fyhn M, Hafting T, Treves A et al. Hippocampal remapping and grid realignment in entorhinal cortex. Nature 2007;446:190–4.

12. Gardner RJ, Hermansen E, Pachitariu M et al. Toroidal topology of population activity in grid cells. Nature 2022;602:123–8.

13. Gardner RJ, Lu L, Wernle T et al. Correlation structure of grid cells is preserved during sleep. Nat Neurosci 2019;22:598–608.

14. Garvert MM, Dolan RJ, Behrens TE. A map of abstract relational knowledge in the human hippocampal–entorhinal cortex. elife 2017;6:e17086.

15. Griffanti L, Salimi-Khorshidi G, Beckmann CF et al. ICA-based artefact removal and accelerated fMRI acquisition for improved resting state network imaging. NeuroImage 2014;95:232–47.

16. Grill-Spector K, Henson R, Martin A. Repetition and the brain: neural models of stimulus-specific effects. Trends Cogn Sci 2006;10:14–23.

17. Gu Yi, Lewallen S, Kinkhabwala A, Domnisoru C, Yoon K, Gauthier JL, Fiete IR, and Tank DW. Cell 2018; 175, 735–750.

18. Hahamy A, Behrens TE. Measuring the spatial scale of brain representations. 2019 Conference on Cognitive Computational Neuroscience. 2019, 2019– 1174.

19. Haxby JV, Gobbini MI, Furey ML et al. Distributed and overlapping representations of faces and objects in ventral temporal cortex. Science 2001;293:2425–30.

20. Kemp C, Tenenbaum JB. The discovery of structural form. Proc Natl Acad Sci 2008;105:10687–92.

21. Klein-Flügge MC, Bongioanni A, Rushworth MFS. Medial and orbital frontal cortex in decision-making and flexible behavior. Neuron 2022;110:2743–70.

22. Klein-Flügge MC, Wittmann MK, Shpektor A et al. Multiple associative structures created by reinforcement and incidental statistical learning mechanisms. Nat Commun 2019;10:4835.

23. Kriegeskorte N, Mur M, Bandettini P. Representational similarity analysis - connecting the branches of systems neuroscience. Front Syst Neurosci 2008;2.

24. Mark S, Moran R, Parr T et al. Transferring structural knowledge across cognitive maps in humans and models. Nat Commun 2020;11:4783.

25. Nichols TE, Holmes AP. Nonparametric permutation tests for functional neuroimaging: A primer with examples. Hum Brain Mapp 2002;15:1–25.

26. Nili H, Wingfield C, Walther A et al. A Toolbox for Representational Similarity Analysis. PLOS Comput Biol 2014;10:e1003553.

27. Park SA, Miller DS, Nili H et al. Map Making: Constructing, Combining, and Inferring on Abstract Cognitive Maps. Neuron 2020;107:1226–1238.e8.

28. Salimi-Khorshidi G, Douaud G, Beckmann CF et al. Automatic denoising of functional MRI data: Combining independent component analysis and hierarchical fusion of classifiers. NeuroImage 2014;90:449–68.

29. Samborska V, Butler JL, Walton ME et al. Complementary task representations in hippocampus and prefrontal cortex for generalizing the structure of problems. Nat Neurosci 2022;25:1314–26.

30. Schuck NW, Cai MB, Wilson RC et al. Human Orbitofrontal Cortex Represents a Cognitive Map of State Space. Neuron 2016;91:1402–12.

31. Smith SM, Jenkinson M, Woolrich MW, Beckmann CF, Behrens TEJ, Johansen-Berg H, Bannister PR, De Luca M, Drobnjak I, Flitney DE, Niazy RK, Saunders J, Vickers J, Zhang Y, De Stefano N, Brady JM, Matthews PM. Advances in functional and structural MR image analysis and implementation as FSL. NeuroImage 2004; 23: S208–S219.

32. Tolman EC. Cognitive maps in rats and men. Psychol Rev 1948;55:189–208.

33. Trettel SG, Trimper JB, Hwaun E, Fiete IR, Colgin LL. Grid cell co-activity patterns during sleep reflect spatial overlap of grid fields during active behaviors. Nature Neuroscience 2019; 22: 609–617.

34. Waaga T, Agmon H, Normand VA et al. Grid-cell modules remain coordinated when neural activity is dissociated from external sensory cues. Neuron 2022;110:1843–1856.e6.

35. Whittington JC, McCaffary D, Bakermans JJ et al. How to build a cognitive map. Nat Neurosci 2022;25:1257–72.

36. Whittington JC, Muller TH, Mark S et al. The Tolman-Eichenbaum machine: unifying space and relational memory through generalization in the hippocampal formation. Cell 2020;183:1249–63.

37. Wilson RC, Takahashi YK, Schoenbaum G et al. Orbitofrontal Cortex as a Cognitive Map of Task Space. Neuron 2014;81:267–79.

38. Winkler AM, Ridgway GR, Webster MA et al. Permutation inference for the general linear model. NeuroImage 2014;92:381–97.

39. Woolrich M W, Ripley BD, Brady M, & Smith, SM. (2001). Temporal Autocorrelation in Univariate Linear Modeling of FMRI Data. NeuroImage, 14(6), 1370–1386

40. Woolrich MW, Behrens TEJ, Beckmann CF, Jenkinson M, Smith SM. Multilevel linear modelling for FMRI group analysis using Bayesian inference. NeuroImage 2004; 21: 1732–1747.

41. Xie J, Padoa-Schioppa C. Neuronal remapping and circuit persistence in economic decisions. Nat Neurosci 2016;19:855–61.

42. Yoon K, Buice MA, Barry C et al. Specific evidence of low-dimensional continuous attractor dynamics in grid cells. Nat Neurosci 2013;16:1077–84.

