## Supplementary information for "Flexible neural representations of abstract structural knowledge in the human Entorhinal Cortex"

*: Equal contribution

### Grid and place cells analysis

#### *Grid cells of all 3 animals generalise over the two different environments*

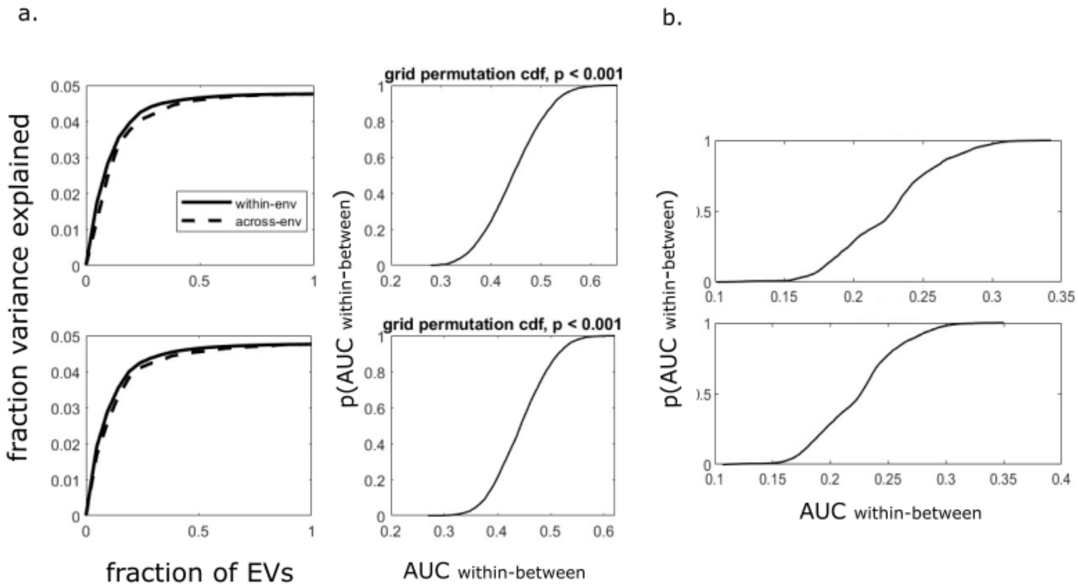


*Figure S1 subspace generalization: grid cells generalise over different environments*

a. *Left: Grid cells subspace generalization graph for 2 more mice (averaged over projection on environment one and two). The lower and upper plots share the same legend. Right: permutation distributions of grid cells (number of permutations = 5000). We permuted the cells in the activity matrix (cells X bins) while the animal forages in one environment, then projected on the EVs from the activity matrix while the animal forages in the other environment and calculated the AUC (and vice versa). We then calculated the difference in AUC between the within environment AUC and the AUC resulting from the permutation and calculated the cdf.*

   *Using these distributions, we can conclude that the difference in AUC of within and between environments is smaller than expected by random projections. p<0.001 for both animals.*

b. *The place cells' sampling distribution of the difference in AUC of within and between arenas (number of bootstraps = 1000, for more details see methods, permutation test 2). We treat these distributions as our NULL distributions to answer the question whether grid cells' AUCwithin-between is smaller than place cells' AUCwithin-between. i.e. do grid cells generalise significantly better than place cells. Upper plot: 14 cells are being sampled, Lower plot: 21 cells are being sampled (to match the number of grid cells that were recorded within each animal), here again the calculation repeated twice, each time we project on one of the environments and then averaged the results.*

***Place cells single mouse analysis:***

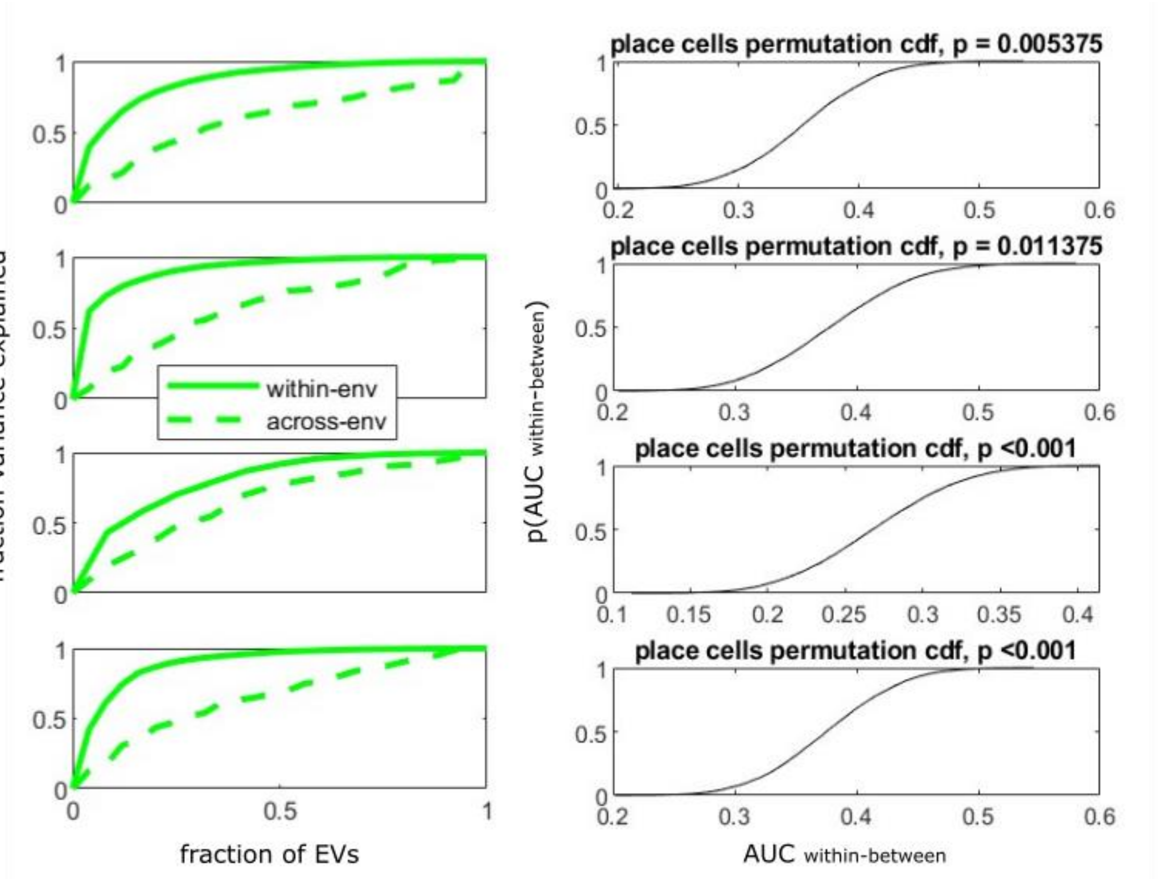


*Figure S2. subspace generalization of place cells*

*Left: subspace generalization plots for the different mice. Solid: within environment; dotted: across environments. Right: the corresponding permutation distribution within the corresponding mouse place cells. Cells were permuted before projection (as before). P-values correspond to the significance of the difference in the AUC for within and across environments under this perturbation distribution.*

In all animals the difference in AUC is smaller than expected by chance, though the effect is smaller than for grid cells. This contradicts the cartoon picture of orthogonal hippocampal representations between environments, but is consistent with models of hippocampal remapping (TEM, (Whittington *et al.* 2020)) and a number of recent empirical findings arguing that hippocampal remapping is non-random (Liu, Sibille and Dragoi 2021; Samborska *et al.* 2022; Tanni, De Cothi and Barry 2022).

**From cells to voxels**

We do not know a-priori how the cells within a module are distributed into voxels. In the main manuscript we segregate each module into four voxels, in Figure S3a, we show that we get similar results if we segregate the module into two voxels. Here, the cells are grouped into voxels according to the rhombus' diagonal. We next examine how the segregation of each module into a different number of voxels influence subspace-generalization score. Here, each voxel is composed of the sum over the cells' activity and noise. The signal to noise ratio (SNR) decreases as the number of voxels increases (Figure S3b) because the number of grid cells within a voxel decreases. The SNR is much larger for voxels with cells that are sampled according to phase compared to randomly (Figure S3b). Therefore, random sampling of cells to voxels leads to subspace generalization score within chance level (Figure S3b). We assume here no correlation between the noise across voxels, which may be unrealistic, full investigation of the noise effect is out the scope of this paper.

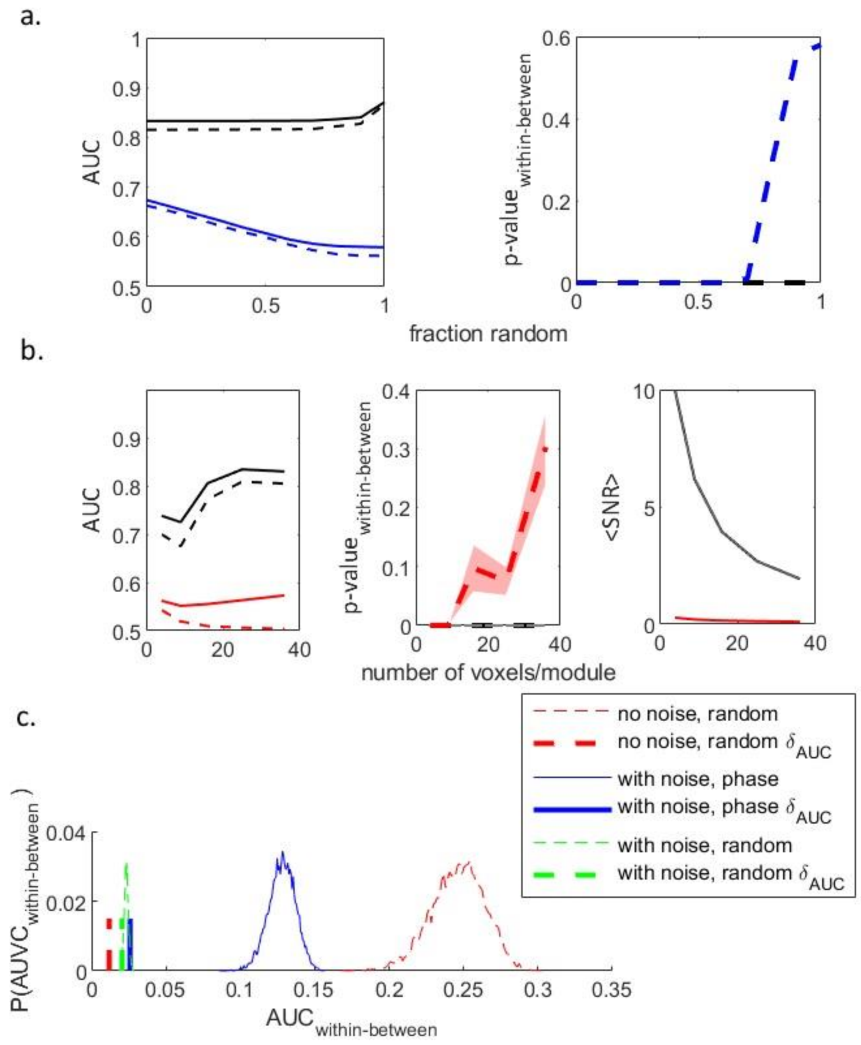

*Figure S3: Dependency of subject generalization on number of voxels per module*
*a. Subspace generalization plot for voxels of the 4 modules, each module is segregated into two voxels. as a function of the ratio of randomly sampled cells. Black – no noise, blue – with noise. left: AUC (solid – within dash – across environments). right: p-value of the effect according to the permutation distribution (see methods, shaded area- standard error of the mean)*
*b. Subspace generalization plot for voxels of the 4 modules as a function of the number of voxels per module. noise std = 50. black – sampled according to phase, red – sampled randomly.*
*c. Left: AUC (solid – within dash – across environments). Middle: p-value of the effect according to the permutation distribution (see methods, shaded area- standard error of the mean). Right: SNR as a function of number of voxels per module.* $SNR \equiv \frac{std(signal)}{std(noise)}$ *were the std(signal) is the averaged standard deviation over the voxels' activity map across the environment. Permutation distributions (the Null distribution). The shift of the distributions to the left following the introducing of noise / random sampling explains the increase in p-values even though the difference in AUC is still small.*

**Subspace Generalisation vs correlating the Neuron-by-Neuron correlation matrices:**

In the following section, we compare Subspace Generalisation as defined in our manuscript to a related and somewhat simpler method for looking at how neurons/voxels covary across tasks: correlating across tasks the (upper triangle of the) Neuron-by-Neuron correlation matrices. We show that Subspace Generalisation emphasises the low dimensional characteristic of the representation and under reasonable conditions should be more robust to spatial noise correlations. Finally, we discuss the reasons for preferring to use the covariance matrix rather than correlation matrix in the calculations of either method.

We can write the correlation between the covariance matrices, Cα, Cβ, as:

$$\rho = \frac{\mathrm{tr}(C_\alpha C_\beta)}{\sqrt{\mathrm{tr}(C_\alpha)\mathrm{tr}(C_\beta)}} = \frac{\sum_q^N U_\alpha^q \,(C_\alpha C_\beta) {U_\alpha^q}^T}{\sqrt{\mathrm{tr}(C_\alpha)\mathrm{tr}(C_\beta)}} = \frac{\sum_q^N \lambda_\alpha^q (U_\alpha^q \, C_\beta {U_\alpha^q}^T)}{\sqrt{\mathrm{tr}(C_\alpha)\mathrm{tr}(C_\beta)}}$$

Because covariance matrices are symmetric we can write the correlation between the upper triangular elements of these matrices as:

$$\rho = \frac{0.5}{\sqrt{\mathrm{tr}(C_\alpha)\mathrm{tr}(C_\beta)}}\left[\mathrm{tr}(C_\alpha C_\beta) - \sum_i C_{ii}^\alpha \, C_{ii}^\beta\right] = 0.5\left[\frac{\sum_q^N \lambda_\alpha^q (U_\alpha^q \, C_\beta {U_\alpha^q}^T)}{\sqrt{\mathrm{tr}(C_\alpha)\mathrm{tr}(C_\beta)}} - \sum_i C_{ii}^\alpha \, C_{ii}^\beta\right]$$

while the AUC in our method is calculated as:

$$\mathrm{AUC} = \frac{\sum_q^N (N - q + 1)(U_\alpha^q \, C_\beta {U_\alpha^q}^T)}{\mathrm{tr}(C_\beta)}$$

We can see that the differences between the two measures are (1) the scaling of each component in the summation (the projection of the matrix on the corresponding eigenvector) (2) the effect of the diagonal elements of the matrix.

Emphasis on low-dimensional representation: We can look at the numerator of the two measures and define $f_\alpha^q \equiv U_\alpha^q C_\beta U_\alpha^{q^T}$ then the numerator becomes:

$$\rho_{numerator} = \sum_q^N \lambda_\alpha^q f_\alpha^q$$

which can be thought of as the covariance between the eigenvector of one matrix and the projection (f), and

$$AUC_{numerator} = \sum_q^N (N - q + 1)\, f_\alpha^q$$

which can be thought of as a covariance of the projection f with a linear decreasing function. This emphasizes the requirement for decreased variance explained as a function of eigenvectors and therefore emphasizes the requirement for low dimensional representation.

The diagonal elements of the covariance matrix give us the variance within each voxel/neuron along the different tasks' states, therefore is larger in areas that encode the states within the task and the signal is distributed across voxels.

Robustness to strong spatial noise: in the fMRI analysis we look at the contrast between 2 conditions, one corresponds to our relevant conditions to compare (here, same structure) and the other corresponds to the control:

The difference in the correlation coefficient can be therefore written as:

$$\Delta\rho = \frac{1}{\sqrt{tr(C_\beta)}} \left( \frac{\sum_q^N \lambda_\alpha^q \left(U_\alpha^q C_\beta U_\alpha^{q^T}\right)}{\sqrt{tr(C_\alpha)}} - \frac{\sum_q^N \lambda_r^q \left(U_r^q C_\beta U_r^{q^T}\right)}{\sqrt{tr(C_r)}} \right)$$

while the difference in AUC can be written as:

$$\Delta AUC = \frac{\sum_q^N (N - q + 1) \cdot \left[\left(U_\alpha^q C_\beta U_\alpha^{q^T}\right) - \left(U_r^q C_\beta U_r^{q^T}\right)\right]}{tr(C_\beta)}$$

From the above we can see that if there is a strong noise correlation, and it is orthogonal to the direction of the signal variance, it will be cancelled out in our method (corresponding to the first eigenvector), while this is not necessarily the case in the difference between correlations (that requires not only the same ordinal position but the exact fraction of the corresponding eigenvalue and the sqrt of the trace). Therefore, in the case of strong spatial noise correlation our method is beneficial. Note that if there is strong spatial noise correlation in one condition and not the other it will damage both methods but because our study controls inputs that are not related to the structure this seems unlikely. But please pay attention that our matrices $C\boldsymbol{\alpha},\boldsymbol{\beta}$ are composed of covariance between the betas and not the pure voxels (see our response to the previous reviewer comment). Further research is needed in order to fully characterize which method is more beneficial and under which conditions.

Using correlation or covariance matrix to calculate the eigenvectors (which is the same question as whether we should standardize the data or not) is an interesting question that requires more thought and research. In our paper we indeed use the covariance matrix that takes into account the scale of the data (voxel's beta/neuron firing rate). Using the covariance matrix indeed allows us to interpret our results as the direction of variance explained. But, it is a reasonable claim that the important measure to define a cell assembly (that may group into voxels) is the correlation between the neurons/voxels and not their covariation. Please note that in this case, the sum over the diagonal elements that is included in subspace generalization but ignored while correlating the upper triangular elements, is just the dimension of the correlation matrix and therefore does not depend on the distribution of the data variance along the different dimension. Nevertheless, because here our variables are the betas from the first level GLM, it makes sense to give less weights, within a search light, on voxels with smaller betas which probably have very small SNR and that also means that their response to our stimuli is low.

<u>**Inferring community structure**</u>

**Learning a community structure: Participants prefer to choose a connecting node even though it is the wrong answer**

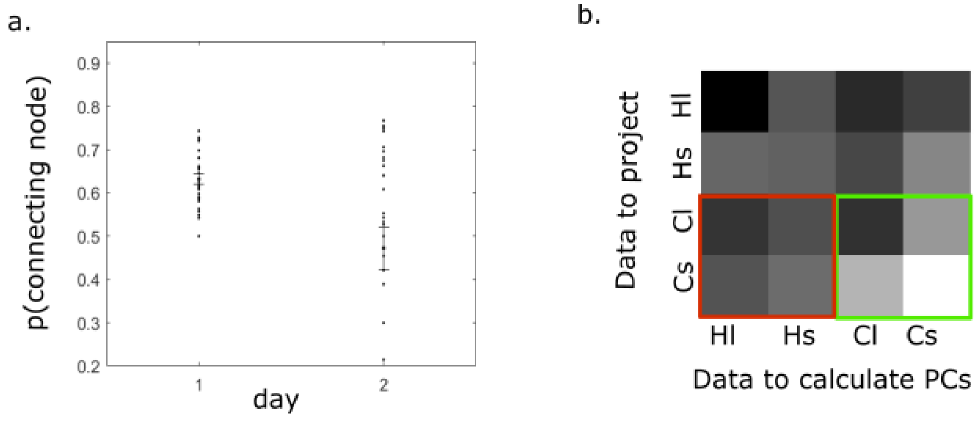

*Figure S4: learning a community structure.*
*a. During the first community structure training day, during the navigation task, participants preferred to choose connecting nodes even though it was the wrong answer.*
*b. Subspace generalization matrix in the PFC (ROI taken from (Baram et al. 2021), 125 voxels around the peak)*

During the first day navigation task (see Figure 3F), when participants had to choose between a picture that is a connecting node or a non-connecting node picture, they preferred to choose a connecting node picture even though it increases the number of steps to the target (Figure S4a, one sided t-test against chance level (50%) $t(27)= 10.8$, $p<<0.001$). This result suggests that participants used the community structure graph to inform their behavior, i.e. the inference of the structure leads to a particular behavioral policy that fits to that particular structure (see also (Mark *et al.* 2020)). This was not true during the second day, when participants' knowledge of the graph improved; participants chose wrong connecting nodes during the first training day significantly more than during the second training day ($t(54) = 3.27$, $p < 0.01$).

**Flexible representation of structural knowledge in PFC**

There is evidence of grid-like activity patterns in mPFC as well as EC while humans perform virtual navigation tasks (Doeller, Barry and Burgess 2010). The parallels between spatial and non-spatial tasks representations suggest that structural representation should be found in PFC as well, though it is still not clear what are the conditions that activate EC, PFC or both. Indeed, hexagonal activity pattern was also observed in PFC when humans navigated abstract 2D Euclidian space and when monkeys had to infer value-based decisions in non-spatial task (Constantinescu, O'Reilly and Behrens 2016; Bongioanni *et al.* 2021). In our experiment, using the method that was described above, we could not find indications for PFC structural representation of hexagonal structure. One option is that PFC exploits structural representation when structural dependent decisions should be taken to gain a reward (Baram *et al.* 2021; Samborska *et al.* 2022).

In our experiment there was no explicit reward. Nevertheless, navigating on a community structured graph may result in internal reward; getting trapped inside a community is frustrating, therefore escaping from a community is satisfying. Further, navigation on community structured graphs requires the exploitation of a specific behavioral policy that is structure dependent (Figure S3, Mark *et al.* 2020). This raises the possibility that a flexible and abstract representation of community structure might exist in mPFC.
When humans had to make decisions in order to gain a reward while exploiting structural knowledge, previous study revealed a structure dependent activation in mPFC (Baram *et al.* 2021). We therefore chose the ROI from Baram et al. to check for structural representation in our task. We have applied the same method as before on voxels in this ROI and checked for generalization of community structure knowledge. We indeed found that subspace generalization for community structure encoding in this ROI is significantly larger within community structure than between structure (i.e the contrast: [ClCl + ClCs + CsCl + CsCs] - [ClHl + ClHs + CsHl + CsHs], coordinate: [-4,44,-20], for this voxel $t(27) = 1.8$, $p<0.05$, for the average of 125 voxels around the peak: $t(27) = 1.79$, $p < 0.05$, Figure S4b), though we note that the

effect is weak. Our results tentatively suggest that mPFC represents the knowledge of community structured graphs abstractly and flexibly, but more experiments should be done to reinforce this conclusion.

Sequences of three images in the graph

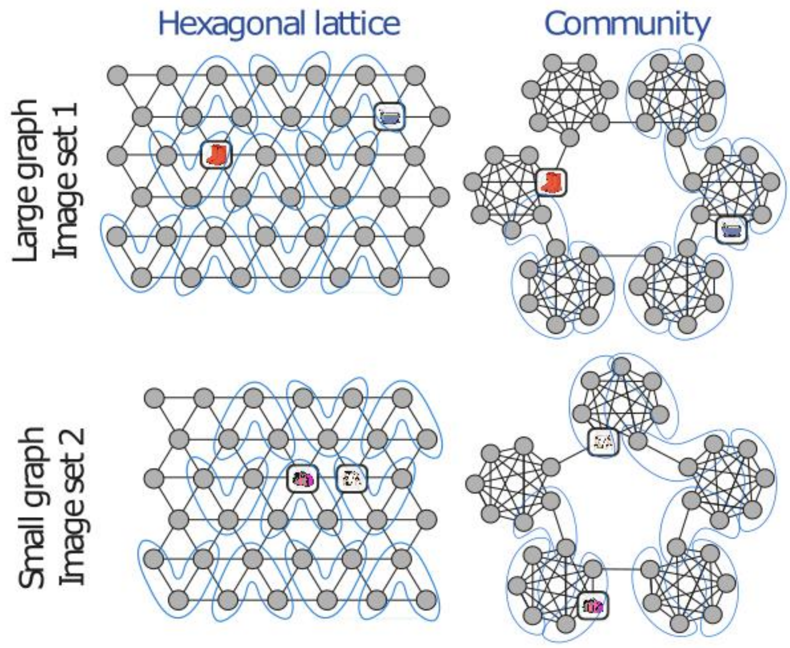


*Figure S5: Partition of the graphs into three images sequences*.

Behavior during the fMRI session

To encourage attention to the sequences during the fMRI session, in 12.5% of trials the sequence was followed by a single image ("catch trial" in Figure 4c), and participants had to indicate whether it was associated with the last image in the sequence. Participants answered these questions significantly better than chance (t-test, $p<<0.001$, $t[27]_{hex}=11.3$, $t[27]_{cluster}=10.6$), for both types of structures, indicating that they: 1) remember the associations 2) recognize the correct graph 3) maintain the correct representation during the block. On each block we calculated the fraction of correct answers and averaged the fractions over all blocks within participant.
At the beginning of the fifth day, we asked participants whether they could describe how the images are associated, and whether they found any difference between the associations in the pictures sets that were learnt during the first two days (i.e. the hexagonal graphs) and the sets that were learnt during the third and fourth day (the community graphs); 26 out of 28 participants mentioned verbally that the pictures in the sets of the last two days were segregated into groups (i.e. community structure). In the scanner, for these 26 participants, we

asked at the end of each fMRI block whether the block's set of pictures contained groups. For the other two participants, we asked whether the set belonged to the first two training days. Below (Figure S6, right), we summarize participants responses. Participants answered correctly which picture set they are currently playing with, significantly better than chance for both structures (t-test, $p<0.001$ for both structures, $t[27]_{hex} = 3.8$, $t[27]_{cluster} = 9.96$).

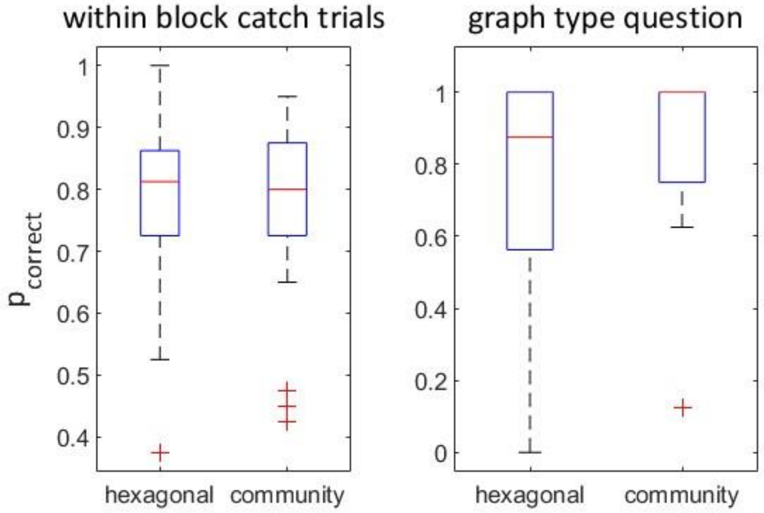


*Figure S6: behavior during the fMRI session.*

*a. Participants were able to detect whether the image in question can indeed follow the current three picture sequence significantly better than chance (t-test, p<<0.001, t[27]hex=11.3, t[27]cluster=10.6)*

*b. Fraction of correct answers to the recognition question at the end of each block. (t-test, p<0.001 for both structures,* $t[27]_{hex} = 3.8$, $t[27]_{cluster} = 9.96$*). Error bars denote SEMs.*

<u>references</u>


Baram AB, Muller TH, Nili H *et al.* Entorhinal and ventromedial prefrontal cortices abstract and generalize the structure of reinforcement learning problems. *Neuron* 2021;**109**:713–23.

Bongioanni A, Folloni D, Verhagen L *et al.* Activation and disruption of a neural mechanism for novel choice in monkeys. *Nature* 2021;**591**:270–4.

Constantinescu AO, O'Reilly JX, Behrens TE. Organizing conceptual knowledge in humans with a gridlike code. *Science* 2016;**352**:1464–8.

Doeller CF, Barry C, Burgess N. Evidence for grid cells in a human memory network. *Nature* 2010;**463**:657–61.

Liu K, Sibille J, Dragoi G. Orientation selectivity enhances context generalization and generative predictive coding in the hippocampus. *Neuron* 2021;**109**:3688-3698.e6.

Mark S, Moran R, Parr T *et al.* Transferring structural knowledge across cognitive maps in humans and models. *Nat Commun* 2020;**11**:4783.

Samborska V, Butler JL, Walton ME *et al.* Complementary task representations in hippocampus and prefrontal cortex for generalizing the structure of problems. *Nat Neurosci* 2022;**25**:1314–26.

Tanni S, De Cothi W, Barry C. State transitions in the statistically stable place cell population correspond to rate of perceptual change. *Curr Biol* 2022;**32**:3505–14.

Whittington JC, Muller TH, Mark S *et al.* The Tolman-Eichenbaum machine: unifying space and relational memory through generalization in the hippocampal formation. *Cell* 2020;**183**:1249–63.
